# Off-target mapping enhances selectivity of machine learning-predicted CK2 inhibitors

**DOI:** 10.1101/2025.11.12.688093

**Authors:** Huiyan Ying, Weikaixin Kong, Aron Schulman, Nikola Panajotovikj, Mirva Pääkkönen, Tanisha Malpani, Ziaurrehman Tanoli, Otto Kauko, Jordi Mestres, Tero Aittokallio, Mitro Miihkinen

**Affiliations:** Institute for Molecular Medicine Finland (FIMM), HiLIFE, University of Helsinki, Helsinki 00290, Finland; iCAN Digital Precision Cancer Medicine Flagship, University of Helsinki and Helsinki University Hospital, Helsinki 00290, Finland; Chemotargets SL, Parc Cientific de Barcelona, Baldiri Reixac 4 (TR-03), 08028 Barcelona, Catalonia, Spain; Institut de Quimica Computacional i Catalisi, Facultat de Ciencies, Universitat de Girona, Maria Aurelia Capmany 69, 17003 Girona, Catalonia, Spain; Turku Bioscience Centre, University of Turku and Åbo Akademi University, 20520 Turku, Finland; Institute for Cancer Research, Department of Cancer Genetics, Oslo University Hospital, Oslo 0310, Norway; Oslo Centre for Biostatistics and Epidemiology (OCBE), Faculty of Medicine, University of Oslo, Oslo 0372, Norway

## Abstract

A key challenge in drug development is identification of druggable targets, the modulation of which attenuates disease progression, while avoiding inhibition of proteins that lead to dose-limiting toxicities. Here, we investigate a drug target casein kinase 2 (CK2) - a serine/threonine kinase implicated in cancer, for which existing inhibitors have so far failed in clinical trials. Using molecular and pharmacoepidemiology approaches, we show that small molecules targeting cyclin-dependent kinase (CDK) family members CDK1/2/7/9, including the existing CK2 inhibitors, have a higher risk to induce adverse effects or fail in clinical trials. Based on this finding, we establish a machine learning (ML) assisted discovery pipeline to redesign more specific and allosteric lead compounds against CK2, with a more selective on-target binding and favourable off-target profile. Importantly, we show that such selective design is possible when standard molecular docking and ML algorithms are combined with an error prediction model. In conclusion, our study reports a simple yet efficient ML-powered drug discovery pipeline and novel submicromolar inhibitors targeting clinically relevant CK2 kinase with no clinically approved antagonists available. Our prediction pipeline was able to achieve a 90% hit-rate, significantly reducing the need for subsequent wet-lab validation.

## Main

In drug discovery, off-targets refer to proteins whose unintended engagement by a therapeutic agent produces severe adverse or dose-limiting effects, without contributing to the desired therapeutic efficacy. Unlike primary or secondary efficacy targets, off-targets are liabilities embedded within the proteome that constrain therapeutic selectivity. Well-recognized examples include the hERG potassium channel (KCNH2), the blockade of which leads to cardiac arrhythmias, and cytochrome P450 enzymes, where broad inhibition causes drug–drug interactions (Goldwaser et al. 2022; Zhang et al. 2014). Systematic mapping of such off-targets is becoming a cornerstone of modern pharmacology as harmonized data on binding affinities, genetic dependencies and drug-induced adverse reactions become increasingly available (Isigkeit et al. 2024; Ianevski et al. 2024; Tanoli, Fernández-Torras, et al. 2025; Elmore et al. 2025; Giri et al. 2019). Increased awareness of off-target activities enhances early screening cascades and guides medicinal chemistry efforts to reduce off-target binding and inform risk-benefit assessments (Schmidt et al. 2025).

In parallel with avoiding off-targets, drug molecules need to effectively bind proteins driving disease progression or resistance to achieve therapeutic benefit. Whether through drug repurposing or *de novo* molecular design, this is increasingly being done in reliance on artificial intelligence (AI) or machine learning (ML) methods, which enable systematic exploration of the small molecular space (Tibo et al. 2024). While AI/ML holds significant promise for addressing long standing drug discovery challenges, such as target prediction and activity profiling, many applications remain limited by data quality, model generalizability, and biological complexity. Within the kinase inhibitor space, AI/ML methods have begun to show potential for improving predictions of binding affinity and selectivity based on chemical structure and bioactivity data (Schulman et al. 2024; Liu et al. 2023; Cichońska et al. 2021; Theisen et al. 2024). However, selectivity modeling, particularly across closely related kinase families, remains an open challenge. More refined approaches, better datasets, and thoughtful, wider integration into drug discovery workflows are needed to realize the potential of ML in guiding kinase inhibitor design, with improved safety profiles via systematic off-target awareness.

In this work, we explore the landscape of off-targets across CDK family members and cancer cell lines, using drug bioactivity profiles, genetic dependencies, and post-marketing safety data. To improve likelihood for clinical success, we implement ML-driven strategies for the rational design of kinase inhibitors with enhanced selectivity profiles. In particular, we train a structurally assisted prediction pipeline, based on support vector machine and molecular docking, which identified novel small molecules against CK2, a serine/threonine kinase relevant for both solid and hematological cancers (Pizzi et al. 2015; Bulanova et al. 2024). Despite the early reports of CK2’s role in oncogenic signalling (Scaglioni et al. 2006), no clinically approved inhibitors for CK2 exist. We perform validation of two of the high-scoring lead molecules in their ability to selectively inhibit CK2 kinase in comparison to other kinases (their off-target profiles). Using phosphoproteomics and *in vitro* kinase assays, we show that our novel molecules effectively inhibit CK2 signalling pathways.

## Results

### Machine learning-assisted discovery pipeline for CK2 small-molecular inhibitors

Casein kinase 2 (CK2) is a heterotetrameric kinase complex and an attractive drug target, as it has been shown to play a critical role (Pizzi et al. 2015) and over-expressed in multiple prevalent malignancies, including breast cancer (**Figure S1**). Its activity potentiates many oncogenic signalling pathways (Borgo, D’Amore, et al. 2021), while the inhibition of CK2 has been shown to synergize with PARP inhibition or conventional chemotherapy in high-grade serous ovarian carcinoma and triple-negative breast cancer (Bulanova et al. 2024). To discover selective CK2 inhibitors, we built an ML-assisted pipeline that first performs site-independent affinity prediction to prioritize compounds, followed by site-specific molecular docking and experimental validation (**Figure 1A**).

**Figure 1.**
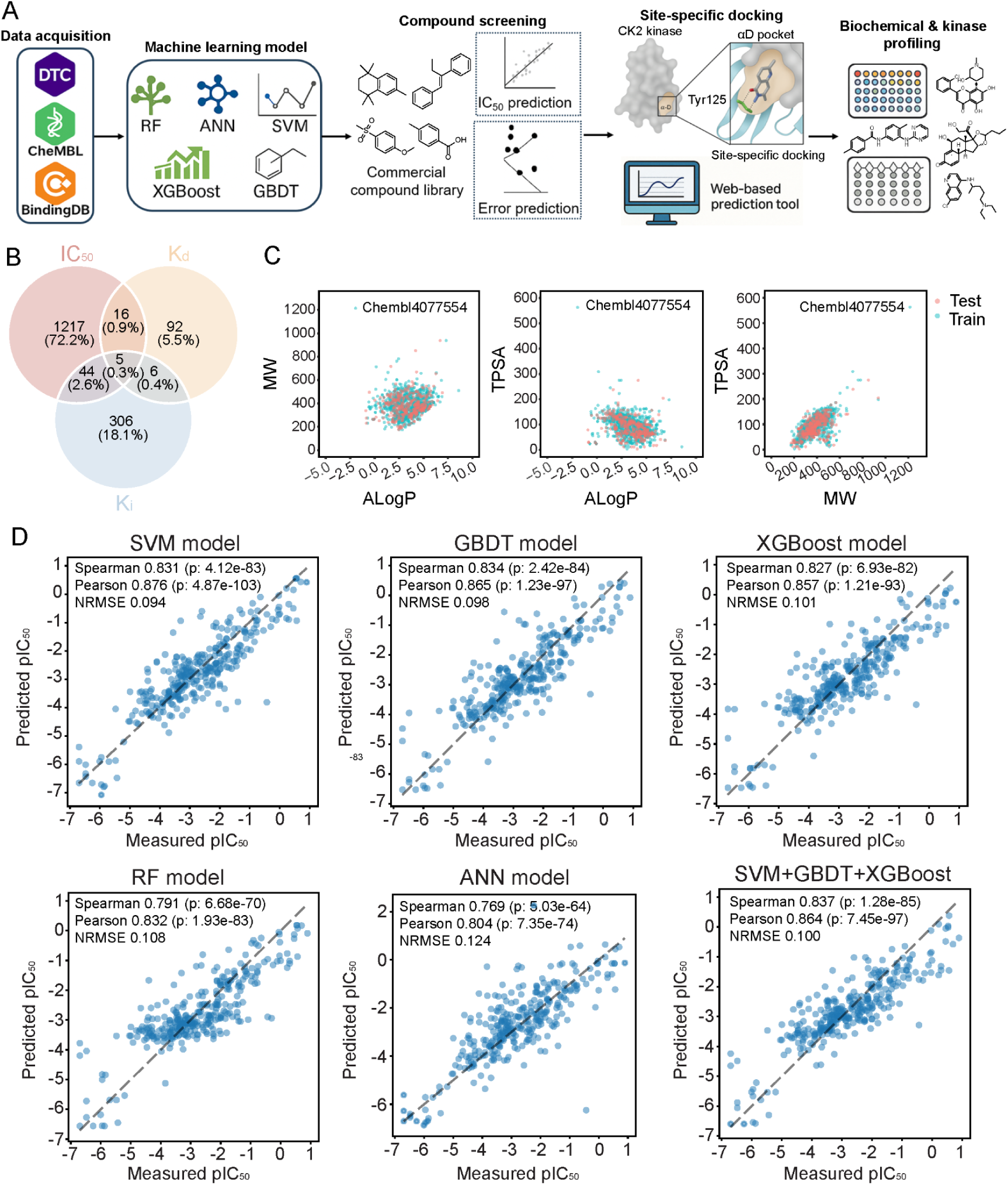
Machine learning-assisted affinity prediction to prioritize lead CK2 inhibitors. **(A)** Schematic illustration of the computational prediction framework followed by experimental validation. **(B)** Venn diagram of compounds with available IC_50_, K_d_, K_i_ bioactivity data against CK2 target. The overlapping portions indicate the compounds with multiple bioactivity types. **(C)** Scatter plots of molecular descriptors in the training and test datasets, including MW vs. ALogP (molecular weight vs lipophilicity), TPSA vs. ALogP (surface polarity vs lipophilicity) and TPSA vs. MW (surface polarity vs. molecular weight). (**D**) Scatter plots between the predicted pIC_50_ and measured pIC_50_ values in the test dataset using various ML models and their ensemble. pIC_50_= -log_10_(IC_50_ ).

The drug/target activity data against CK2 was collected from open databases (**Figure 1A-B**). As shown by multiple prior studies attempting to escalate model complexity, the prospective performance of AI/ML based modelling is dominated by factors such as dataset size, coverage, diversity and quality (Ahlmann-Eltze et al. 2025; Tanoli, Schulman, et al. 2025). Because of this, we placed special emphasis on systematic data-centric optimization including (i) curated and harmonized CK2 activity records across public sources, (ii) standardized structures and assay units/metadata, removed duplicates and reconciled conflicts, (iii) combined activity types (IC_50_, K_i_, K_d_ and their combinations), and (iv) evaluated structural fingerprints alongside chemical-space coverage and similarity-aware splitting to quantify their effects on accuracy and stability. These analyses indicated that IC_50_ labels encoded with a 1024-bit standard fingerprint led to the most reliable performance (**Figure S2-5;** see **Methods**). Accordingly, we used 1,282 non-redundant IC_50_-annotated compounds, represented by the standard fingerprint, for model development under a 3:1 train-test split.

We next assessed the training and test dataset distributions using small molecular descriptors, showing that both datasets encompass a large and diverse range of chemical space. Importantly, the wide chemical space in the training dataset ensured the prediction model has a wide application domain, making it possible to achieve a generalizable accuracy on unknown compounds as well (**Figure 1C**). In addition, we further included molecules with extreme values (e.g. Chembl4077554) outside of Lipinski-Egan-Verber’s laws, as they could contribute to enhancing the model generalizability and accuracy (Cichońska et al. 2021).

To implement a predictive ML model for drug/target affinity prediction, we trained and tested multiple supervised machine learning models from different model families, including random forest (RF), artificial neural network (ANN), support vector machine (SVM), extreme gradient boosting (XGBoost) and gradient boosted decision tree (GBDT). We found that the SVM model had the best performance in terms of both Pearson and normalized root mean square error (NRMSE). For the Spearman coefficient, the GBDT was the top performer, but its relative difference with the SVM model was rather small. In addition, we also ensembled the top 3 models (SVM, GBDT, XGBoost). However, no significant improvement was observed (**Figure 1D**), indicating that the top models make rather similar predictions. In previous studies, SVM has exhibited good performance in the virtual screening (Liang et al. 2020; Han et al. 2008; Jorissen and Gilson 2005), so we selected the SVM model for the subsequent prediction and virtual screening tasks.

To further assess the applicability domain of the bioactivity prediction model and enhance its prediction reliability, we established an error prediction model (EPM) on top of the SVM method and identical parameter settings to those of the primary bioactivity model. The EPM predicts the expected error of the bioactivity prediction model for each compound, allowing exclusion of samples with large predicted errors under different confidence levels (α values). Applying the EPM on top of the SVM model led to significant improvements in the accuracy metrics (**Figure S6,** see **Methods**), demonstrating its potential to refine bioactivity predictions within the model’s applicability domain.

Using our highest performing machine-learning model (SVM model combined with an EPM, we predicted novel CK2 binders out of a large chemical library of 12 M small molecules, while filtering out molecules with poor bioavailability by applying a filtering score based on Lipinski’s rule of five. Out of these molecules, we selected the top 622 molecules (predicted IC_50_<100 nM) and continued with high-throughput docking predictions for the assessment of molecular affinities, binding modes and site-specificity.

### Structure-based prioritisation of novel CK2 hit molecules reveal allosteric inhibitors

Although the SVM model performs well in predicting bioactivity, the models trained merely on CK2 binding data are structurally unaware. Therefore, we next assessed the possible CK2 binding sites for the top 622 molecules predicted to have a high affinity towards the CK2 ɑ-subunit. Interestingly, CK2 ɑ-subunit hosts multiple binding pockets on its surface, enabling potential binding of our predicted small molecules (**Figure 2A**). Between the ATP site, the αD pocket, the CK2α/β interface, and the N-terminal tail, we focused our efforts on the αD pocket which has a lower sequence and structural similarity to other kinases and high potential for allosteric modulation of kinase activity (Brear et al. 2020; Iegre et al. 2021). The CK2α/β interface and the N-terminal were not prioritized since they are shallow and partly solvent exposed, often depending on the β subunit, and having multiple previously reported ligands (Kufareva et al. 2019; Brear et al. 2020; Iegre et al. 2021). Guided by this rationale, we ran two complementary docking schemes: a bitopic protocol that searches for poses contacting the ATP region while extending into the αD pocket, and an αD-only protocol restricted to the αD cavity to capture purely allosteric binders (Brear et al. 2020; Iegre et al. 2021).

**Figure 2.**
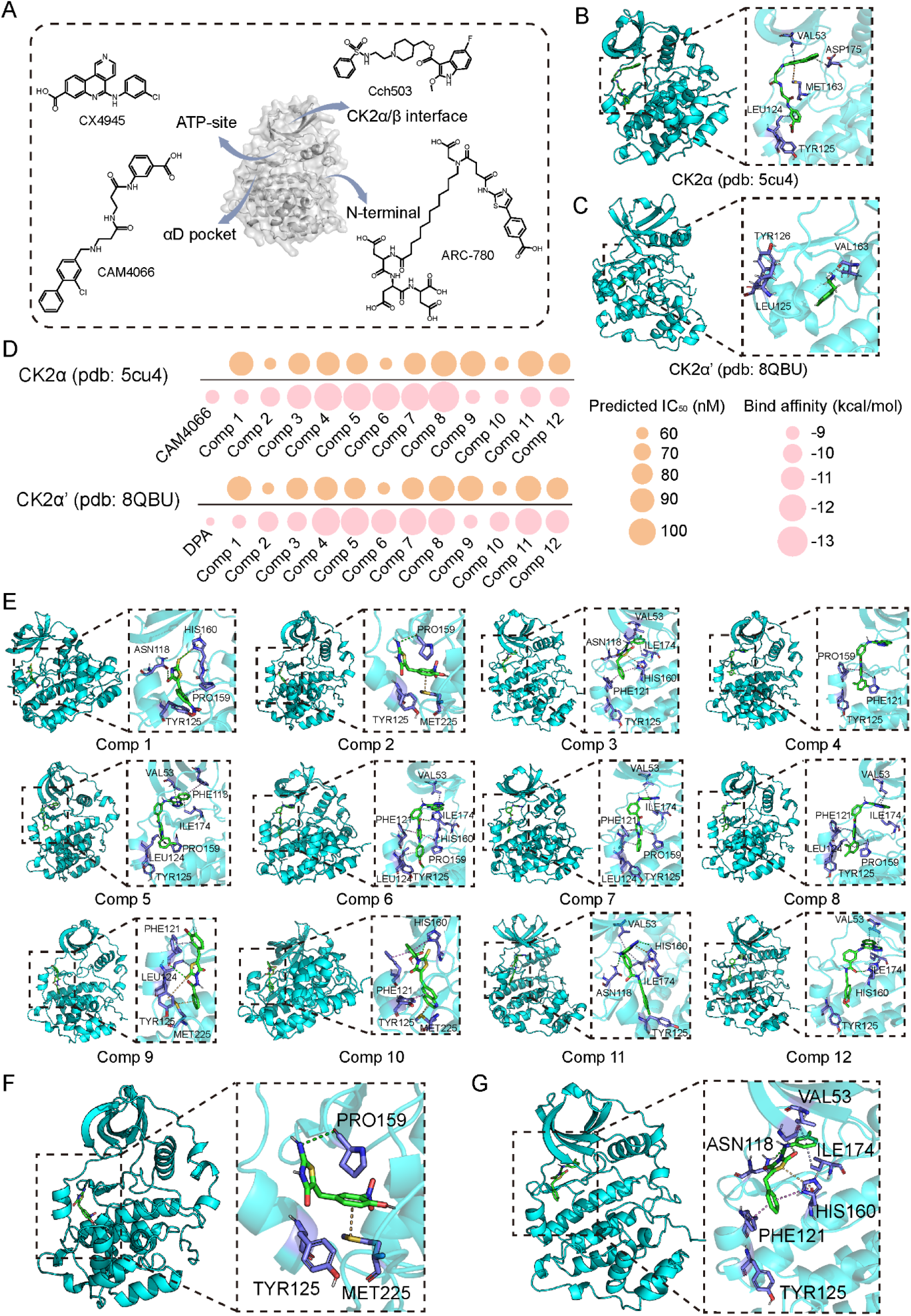
Molecular-docking assisted structure-based prioritisation of lead CK2 inhibitors. (**A**) The structure of CK2 (CK2α unit as an example). Key binding sites and corresponding control compounds are shown (ATP-site/CX4945, αD pocket/CAM4066, CKα/β interface/Cch503, N-termimal/ARC-780). αD pocket as the main docking site to screen the potential inhibitors with two major binding modes with their control compound of (**B**) CAM4066 binds to both ATP-site and αD pocket; (**C**) DPA binds to αD pocket. (**D**) Dot plots illustrate the predicted IC_50_ and binding affinities of the top 12 screened compounds against the αD pocket in both CK2α and CK2α′. All selected compounds exhibit stronger binding (i.e., lower affinity values) compared to the control compound (CAM4066: -9.1 kcal/mol; dichlorophenyl (DPA): -5.5 kcal/mol). (**E**) Conformation of the screened 12 protein-compound structures. Among them, Comp1, Comp2, Comp9, Comp10 mainly bind to αD pocket; Comp3, Comp4, Comp5, Comp6, Comp7, Comp8, Comp11, Comp12 bind to both ATP-site and αD pocket. (**F-G**) Magnification of two of the most promising docking poses, compounds 2 and 3.

To enable high-throughput molecular docking, we took advantage of two previously published crystal structures for CK2ɑ and CK2ɑ’ (**Figure 2B-C**). In both of the cases (ATP-site + αD pocket or αD-pocket docking), we optimized the docking parameters using control compounds, CAM4066 and 3,4-dichloro phenethylamine (DPA), for which the affinity and docking pose to these sites are known. We then ran the docking for the top 622 small-molecules towards CK2alpha or CK2alpha’ and ranked the compounds based on their binding poses and binding affinity as predicted by autodock vina (Trott and Olson 2010). Twelve compounds satisfied one of the two predefined binding modes and showed more favorable docking scores than the control ligands (CAM4066 for CK2α and DPA for CK2α′), and were selected as the potential allosteric CK2 binders. Predicted binding affinities and IC_50_ values are summarized in **Figure 2D**, and representative bound conformations are shown in **Figure 2E**.

Four compounds (Comp1, Comp2, Comp9, and Comp10) were observed binding within the αD pocket, having potential to allosterically inhibit CK2. The remaining eight (Comp3-8, Comp11, and Comp12) adopted bitopic poses that contact the ATP region and extend into the αD pocket. Magnified views of the two most promising docking poses (Comp 2 and 3), are provided in **Figure 2F-G**. Comp2 exemplifies the αD-only mode and forms a hydrogen-bonding network involving Tyr125 and nearby backbone atoms around Pro159, together with aromatic stacking against Met225. Comp3 illustrates the bitopic mode, bridging the ATP hinge region and the αD cavity, with hydrogen-bonding to Asn118, Pi-Pi contacts to Ile174, Val53, His160 and Phe121 and hydrophobic interactions with residues such as Tyr125.

As a conclusion, when using well-harmonized datasets, the SVM machine learning model with parallel error prediction predicts CK2 binders, which can be docked into both ATP or alternative binding pockets, with potential to reveal allosteric inhibitors.

### CDK family kinases are essential off-target proteins associated with clinical attrition

Our ML-assisted discovery pipeline was able to pinpoint multiple small molecules with significant affinity and reasonable docking poses towards CK2. To help selecting the most promising molecules and to maximise the therapeutic window of the lead compounds, we sought to understand the most central off-targets in drug development. Although previous work has suggested frameworks for safety panel design (Schmidt et al. 2025), we performed a more systematic analysis of possible off-targets leveraging the DepMap resource - a cancer centric database that hosts data from whole-genome loss-of-function screens performed for over 1300 cancer cell lines. We hypothesized that genes essential for overall cell viability and survival across the majority of cancer cell lines can be considered as potential off-targets, as their cellular function is likely preserved also in non-malignant cells.

We ranked all recorded genes based on their cell line dependence probability, displaying a right-skewed, heavy-tailed distribution, where 1696 of the genes (9.5%) were classified as highly dependent overall for cellular growth or survival (**Figure 3A**). Analysis of the 1696 most dependent genes across all cell lines identified a plethora of protein groups with a statistically significant overrepresentation. These included gene groups encoding essential machinery for ribosome, spliceosome, proteasome and RNA polymerase (FDR<0.05; **Figure S7A**). Since the most likely off-targets for CK2 targeting molecules are other kinases, due to their structural similarity, we next limited our analysis to cellular kinases only. This analysis highlighted the CDK family of kinases as a key group of kinases necessary for cellular growth or survival (FDR<0.05; **Figure S7B**) Interestingly, none of the existing safety pharmacology panels list CDK family members as toxicity-inducing targets (Schmidt et al. 2025).

**Figure 3.**
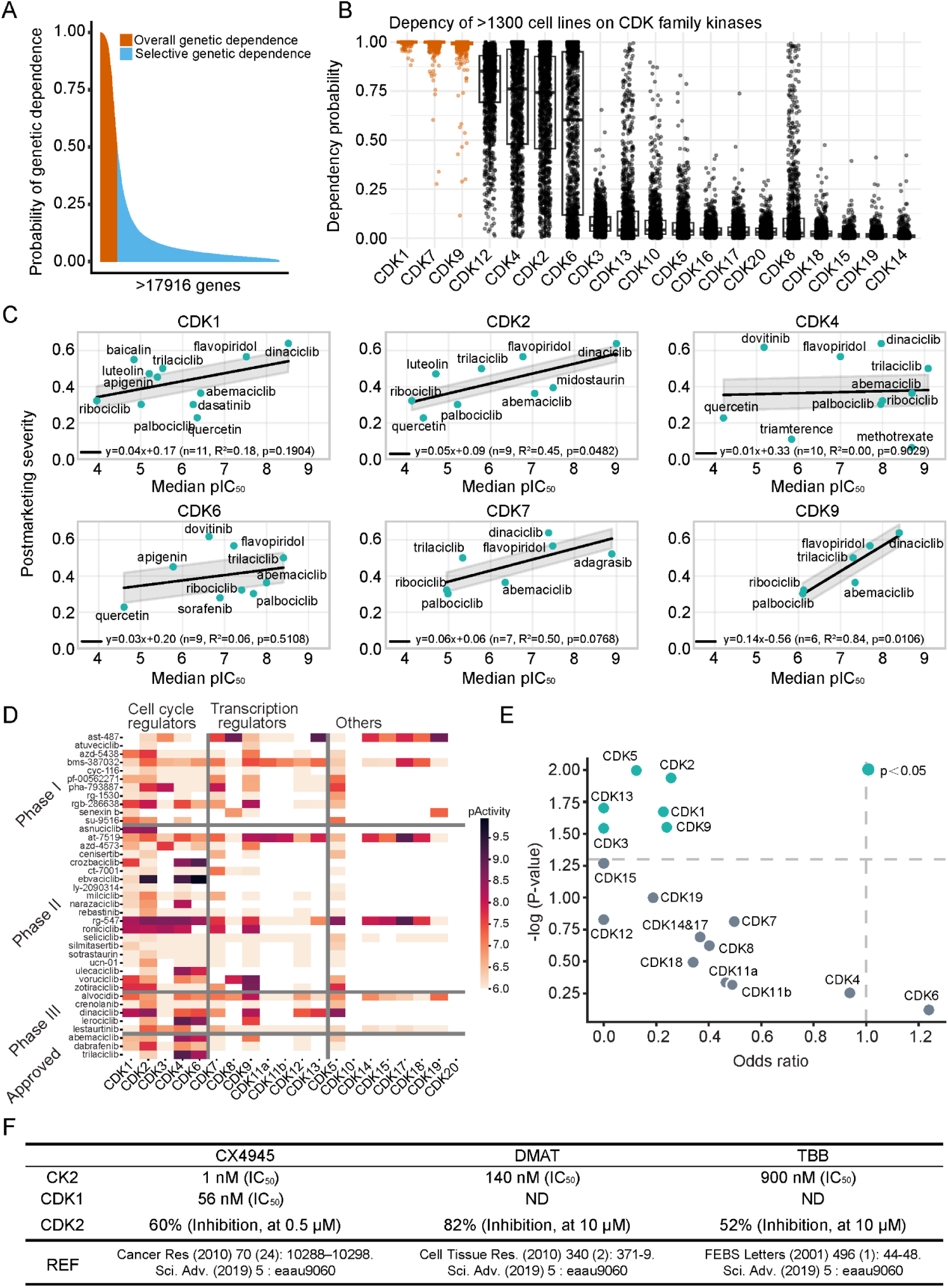
Off-target bioactivity of small molecules against the CDK family contributes to clinical trial attrition. **(A)** Distribution of genetic dependencies across >1300 cancer cell lines (DepMap release 25Q2). Gene dependency probabilities for each gene were averaged over all cell models plotted from high to low. We set the anti-target cut-off to 0.5, which led to 1696 genes defined as highly dependent (anti-targets). (**B**) Dependency probabilities of >1300 cell lines across CDK family members (DepMap release 25Q2). Cell lines are overly dependent on CDK1/7/9. **(C)** Correlation between CDK binding and drugs post-marketing severity score (see **Methods**). (**D**) The bioactivity profiles of selected investigational and approved kinase inhibitors against the CDK family. See **Figure S9** for the complete heatmap. **(E)** Left-tailed Fisher’s exact test was used to assess whether approved drugs show significantly less activity against CDK family members compared to investigational compounds. (**F**) Existing CK2 inhibitors (CX-4945, DMAT and TBB) exhibit off-target activity towards CDK1/2. ND: Not determined.

Based on these findings, we next performed a more detailed genetic dependency analysis, focusing solely on the CDK family of kinases. Interestingly, cells showed variable responses to knockouts of different CDK family members. Here, cell line growth and survival depended on three kinases CDK1/7/9 with a near absolute certainty, and kinases CDK2/4/6/12 showed high dependency scores as well (**Figure 3B**). Since kinases are important targets for multiple small molecules (71 small-molecule kinase inhibitors are currently FDA approved, additional 16 approved by other regulatory agencies) (Attwood et al. 2021), we next explored the safety profiles of clinically approved molecules with activity towards the highlighted CDK family.

Specifically, we assessed whether CDK off-target activity of approved pharmaceuticals is associated with a higher drug toxicity. To understand whether *in vitro* affinities would be associated with more severe side-effects, we first compiled drug safety information from SafetyVista platform (SafetyVista, Chemotargets SL, 2025. https://safety-vista.com/), establishing an extensive database of adverse reactions associated with clinically used pharmaceuticals. We then assessed whether CDK off-target activity negatively impacted the safety profiles of already approved pharmaceuticals. For this we employed a post-marketing drug severity score (see **Methods**), which integrates the percentage of death-related reports and the disproportionality signals of the administered drugs. The severity score ranges from 0 (safer, lower-severity drugs) to 1 (more severe, risk-prone drugs). Our analysis revealed a trend in which off-target activity towards CDK1,2,7 and 9 was associated with more severe drug adverse effects among clinically used pharmaceuticals (**Figure 3C)**. Interestingly, CDK4/6, which is the primary anti-cancer target in multiple malignancies, did not show a similar activity-safety correlation.

After establishing the relationship between CDK targeting and increased drug severity score, we wanted to understand whether bioactivity towards any individual CDK family member would link to increased clinical attrition during clinical testing of potential drug candidates. Using the ChEMBL database, we established a dataset of both pre-clinical and clinical compounds with recorded bioactivity towards the CDK family of kinases (**Figure 3D**). To identify the CDK family members with the highest negative associations with clinical failure, we performed a univariate regression analysis for each CDK family member, evaluating the association between off-target kinase binding and clinical success (**Figure 3E**). Interestingly, activity of a molecule towards 6 out of the 20 CDK kinases had a statistically significant negative association with clinical trial success, suggesting CDK off-target activity towards these molecules contributes to clinical attrition of small molecules during clinical trials. CDK1/2 were among the most statistically significant molecular targets associated with clinical attrition (**Figure 3E**). Out of the CDK family, the only kinase showing slight positive association was CDK6, a known drug target in multiple cancers with many clinically approved molecules binding CDK4/6 as their primary target.

Interestingly, CK2 kinase inhibitor development is one example where molecules targeting the primary target (CK2) have significant off-target effects via the CDK family of kinases and where the previous inhibitors have failed in clinical trials (**Figure 3F**). To gain mechanistic insight into how existing CK2 inhibitors cause toxicities, we integrated data from chemical and toxicity databases, linking small molecular inhibitors with toxicities via their reported CDK family bioactivities (**Figure S8**). This analysis flagged many CDKs, CDK1/2/7 in particular, as relevant off-targets to be considered in preclinical drug discovery, the inhibition of which may lead to systemic toxicities. Notably, currently used safety pharmacology panels, Safety-77 (Brennan et al. 2024) and Safety-44 (Bowes et al. 2012) do not include any CDK kinases in their recommended off-target list (Schmidt et al. 2025).

Based on these results, we argue that previous drug development processes have not adequately steered away from targeting the CDK family of kinases, increasing clinical failure rates. To enable CK2 inhibitor development with enhanced safety profiles and less CDK binding, we next focused on a more refined molecule selection, taking into account both efficacy and toxicity, and aiming for more balanced therapeutic profiles of the molecules targeting CK2.

### Mechanistic assessment of the novel CK2 inhibitors show improved specificity

To validate the ML-model predictions, we next sought to experimentally test the performance of the top 12 predicted molecules, together with 2 positive control molecules and 2 negative molecules. To assess IC_50_ values, we set up an *in vitro* biochemical assay to measure the efficacy of the predicted molecules. This plate-based assay records the amount of ATP used by the kinase, while different compounds can be titered in (see **Methods**). Importantly, 11 of the predicted 12 molecules (discovery success rate >90%) showed a clear inhibition of the target kinase, whereas negative controls did not cause inhibition (**Figure 4A**). Of note, although the positive control molecules CX4945 and SGC-CK2-1 were more efficient in inhibiting CK2 kinase in this reductionist assay, they have so far failed in clinical trials, potentially due to the off target effects and CDK binding. As shown in **Table S2**, CX-4945 displays measurable binding affinity toward multiple CDK family members, including CDK1 (IC_50_ = 56 nM, K_d_ = 3035 nM), CDK2 (IC_50_ = 1800 nM), CDK5 (K_d_ = 514 nM), and CDK16 (K_d_ = 1208 nM). Such broad cross-reactivity across cell-cycle kinases suggests that its cellular effects may not arise solely from CK2 inhibition, but also from off-target CDK blockade, which could contribute to the toxicity and limited clinical tolerability. Since molecules 2 and 3 showed promising inhibition of CK2 (IC_50_ = 156.2 nM and 198.6 nM), we next assessed their off-target binding to other kinases.

**Figure 4.**
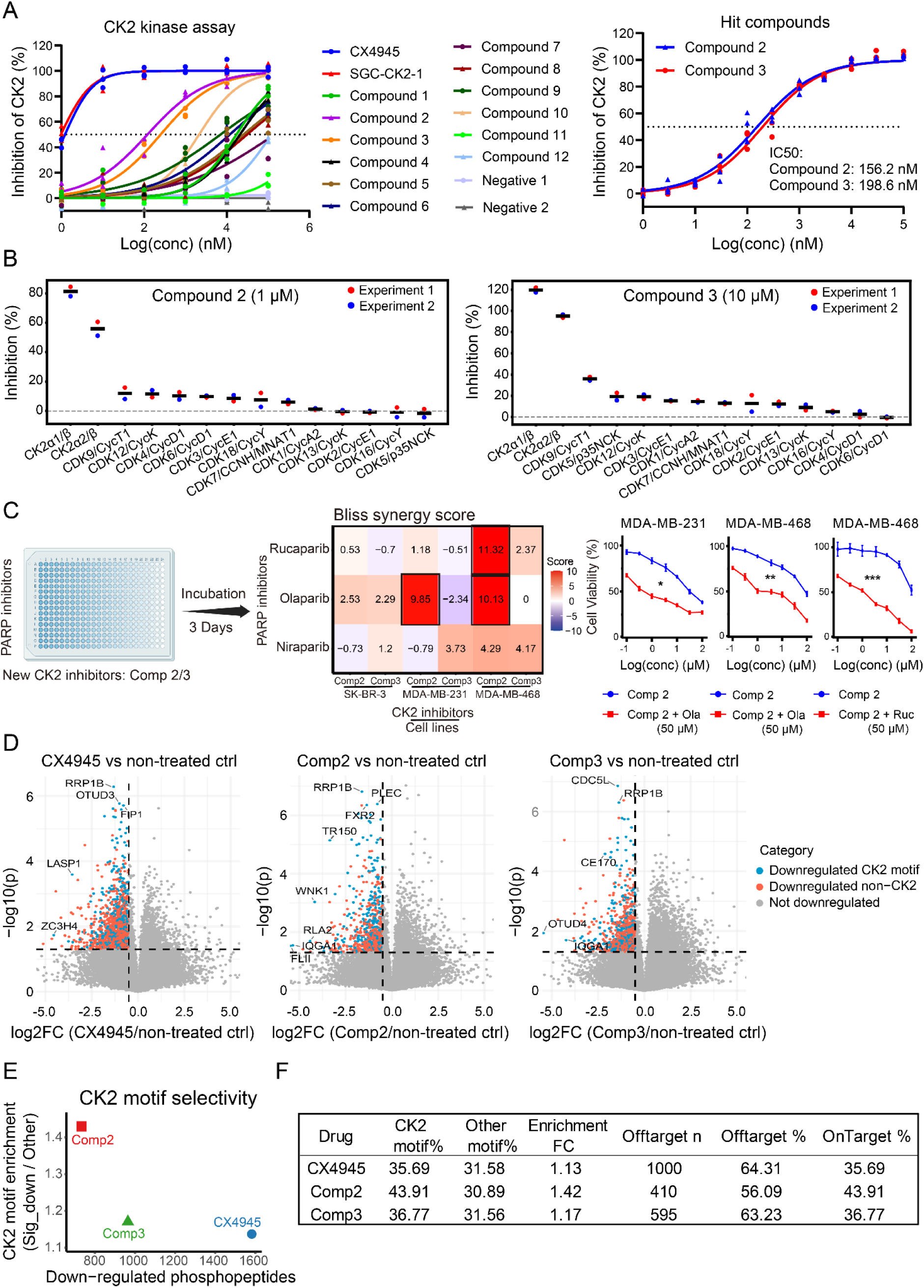
Newly identified CK2 inhibitors are specific to CK2, avoiding off-targets with the toxicity-inducing CDK family. (**A**) CK2 kinase activity assay performed with the 12 prioritized compounds, two positive control (CX4945, SGC-CK2-1), and two negative control (Doxycycline and Ampicillin) at various concentrations (1, 10, 100, 1000, 10000, 100000 nM) (n=3). CK2 kinase assay was re-tested with two hit compounds (Compound 2 and Compound 3, right panel). IC_50_ values were calculated with Graphpad Prism. (**B**) A larger panel of 216 kinases were tested in single-dose activity assay to evaluate the inhibitory effects of Compound 2 (1 μM) and Compound 3 (10 μM) (n=2). Dot plot shows inhibition of CK2 and CDK family kinases. **(C)** Cell lines were treated with a combination of PARPi (Rucaparib, Olaparib and Niraparib) and new CK2 inhibitors (Comp2 and Comp3) in a CTG viability assay. The drug synergies were calculated using SynergyFinder and plotted using the Bliss synergy score. We further tested Comp 2 dose response in MDA-MB-231 and MDA-MB-468 cells in the presence of 50 µM olaparib or 50 µM rucaparib. **(D)** MDA-MB-231 cells were treated for 2 hours using existing (CX4945, 1 μM) or novel (Comp 2/3, 10 μM) CK2 inhibitors and subjected to mass-spectrometry phosphoproteomic analysis (6 technical repeats for each condition). Volcano plots show significantly down-regulated phosphopeptides including downregulated proteins with CK2 motif (S/T-X-X-D/E) (blue) and non-CK2 motif (red) in MDA-MB-231 cells treated for 2 h with CX4945, Comp 2, or Comp 3, and compared with vehicle control. Thresholds used were log2FC ≤ –0.5 and p ≤ 0.05. **(E)** Relationship between global inhibition strength (number of down-regulated phosphopeptides) and CK2 motif selectivity (fold enrichment of motif in down-regulated vs. other phosphopeptides). CX4945 showed the strongest overall suppression but only modest CK2 selectivity, whereas Comp 2 displayed the highest CK2 motif enrichment, suggesting the cleanest CK2-biased inhibition profile. Comp 3 exhibited intermediate suppression and selectivity. **(F)** Summary table of CK2 motif statistics.

To measure the off-target binding and anticipate potential safety liabilities of our top CK2 inhibitors, we tested the bioactivity of Comp2 and 3 against a panel of 216 kinases. Importantly, both of the top molecules were confirmed as strong inhibitors for CK2 with only minor activity towards CDK family members (**Figure 4B**). At a screening concentration of 1 μM, compound 2 exhibited minimal inhibition across the CDK family, with all tested CDK-cyclin complexes showing ≤12% inhibition. In particular, the key cell-cycle regulators such as CDK1/CycA2 (1.28%), CDK2/CycE1 (−0.70%), CDK4/CycD1 (10.25%), and CDK6/CycD1 (9.81%) were largely unaffected, and several CDKs displayed no measurable inhibition (**Table S3**). To further challenge CDK selectivity, compound 3 was profiled at an elevated concentration of 10 μM. Even at this higher dose, compound 3 retained a favorable off-target profile, with most CDK family members exhibiting less than 20% inhibition, including CDK1/CycA2 (14.65%), CDK2/CycE1 (12.36%), CDK3/CycE1 (15.33%), and CDK4/CycD1 (2.57%) (**Table S4**). Moderate inhibition was observed only for CDK9/CycT1 (36.06%) at 10 μM. Interestingly, compound 2 seemed to inhibit the DYRK family of kinases (**Table S3)**, even if the cell line dependency data from DepMap indicates minimal DYRK dependency, when compared to the toxicity-inducing CDK family (**Figure S10**).

Previous studies have shown that CK2 inhibition alone is not enough for killing breast cancer cells, however, its inhibition can synergize with PARP inhibitors or conventional chemotherapy (Bulanova et al. 2024). To assess if our new, more specific CK2 inhibitors synergize with PARP inhibition in breast cancer, we first treated a panel of 8 breast cancer cell lines only with compounds 2 and 3. Congruent to previous results, the cell-killing effect of new CK2 inhibitors was limited, and high concentrations of the compounds were needed to reduce breast cancer cell viabilities (**Figure S11A-B**). To assess drug combination synergy, we next treated cells with combinations of different PARP inhibitors plus CK2 inhibitors Comp2 and 3. In these experiments, the compound 2 synergized with rucaparib and olaparib in most of the tested cell models (**Figure 4C**). Similarly, the compound 2 boosted chemotherapy effectiveness in triple-negative breast cancer models (**Figure 4C**). Only the SK-BK-3 cell line, which is known to be PARP insensitive, did not benefit from the PARPi+CK2i combination therapy (**Figure 4C**).

To assess the inhibitory effects of compounds 2 and 3 on CK2 downstream signalling, we next performed phosphoproteomics assay on cells treated with existing or novel CK2 inhibitors. For the phosphoproteomic analysis, we used the breast cancer MDA-MB-231 cell line, as it is known to express CK2 and it showed synergistic effects in the combination treatment experiments (**Figure 4C**). In these unbiased, multi-replicate experiments, 2 hour CK2 inhibition using existing (CX4945) or novel (Comp2/3) inhibitors did not cause differences in total protein abundance (**Figure S11C**); however, they induced significant changes in protein phosphorylation in all the treated conditions, negatively affecting previously discovered CK2 target proteins such as Farnesoid X receptor (FXR) (**Figure 4D**) (Bilodeau et al. 2017).

Since the CK2 is an acidophilic Ser/Thr kinase and preferentially phosphorylates substrates containing the pS/pT-X-X-D/E motif (Borgo, Cesaro, et al. 2021), we evaluated inhibitor specificity by focusing on peptides that incorporate this characteristic acidic phosphorylation motif. This analysis highlighted Comp 2 as a superior CK2 inhibitor over the previous CX-4945 compound, which showed less efficacy at inhibiting the protein phosphorylation of peptides with the acidophilic motif. In addition, CX-4945 broadly inhibited the phosphorylation of multiple peptides with phosphorylation motifs other than pS/pT-X-X-D/E, suggesting nonspecific cellular effects via off-target kinases such as the CDK family. Compound 3 showed slightly enhanced CK2 motif enrichment, compared to the control inhibitor, but was less efficient than Compound 2, suggesting that this molecule might benefit from further design-test-make cycles (**Figure 4 E-F**). These findings were confirmed using immunoblotting with an antibody recognizing the canonical, phosphorylated CK2 consensus motif. Here, Compounds 2 and 3 inhibited CK2 substrate phosphorylation in all the three tested breast cancer cell lines in a dose-dependent manner, further confirming our findings from the phosphoproteomic analyses (**Figure S11D**).

As a conclusion, we demonstrated that inhibitors from the ML-assisted pipeline are effective on inhibiting CK2 in both *in vitro* biochemical assays and cell-based assays, and show pronounced specificity compared with the tested control kinase inhibitors.

### Structural similarity and scaffold novelty of the new CK2 specific inhibitors

Finally, we wanted to confirm that our discovered compounds are structurally novel and not derivatives of existing inhibitors. To assess scaffold novelty, we first performed a ChEMBL database search, revealing Comp2 as a pre-existing molecule with known affinity towards DYRK1A, and hence confirming findings from our 216-kinase panel screen (**Table S3**). In addition, this molecule has previously been published as a GSK-3β inhibitor (Gandini et al. 2018); however, our experimental validation using the 216-kinase panel showed high bioactivity towards CK2 and DYRK family, with almost no bioactivity towards GSK-3β at 1 μM (**Table S3**). The CK2 inhibitor Comp3 did not exist in these database searches.

We then performed a PCA projection based on ECFP fingerprints, a class of topological fingerprints designed for molecular characterisation (Rogers and Hahn 2010). This analysis revealed that the two new compounds (Comp2 and Comp3) occupied a distinct region of the chemical space, when compared with the ten known CK2 inhibitors (**Fig. 5A**). The established inhibitors clustered according to their chemical series, such as benzimidazoles (TBB, TDB, TBI, DMAT, and TMCB) and indole-based inhibitors (CX-4945, CX-5011, CX-5279), whereas Comp2 and Comp3 formed a separate cluster, indicating chemical distinctness. For quantitative similarity analysis, we used both ECFP4 and ECFP6 fingerprints as they capture structural similarity at different radii, capturing both local atomic environments (ECFP4, radius of 2 bonds) and more global substructures (ECFP6, radius of 3 bonds). These quantitative similarity analyses confirmed the low structural overlap of Comp 2 and 3 with the known CK2 inhibitors (**Fig. 5C**), suggesting low similarity both at the local substructure level (ECFP4) and at more extended, scaffold-relevant molecular environments (ECFP6). Both compounds showed the highest similarity to SGC-CK2-1, yet remained well below the typical threshold (0.6-0.7) for scaffold-level similarity (Zahoránszky-Kőhalmi et al. 2016), suggesting novel chemical frameworks.

**Figure 5.**
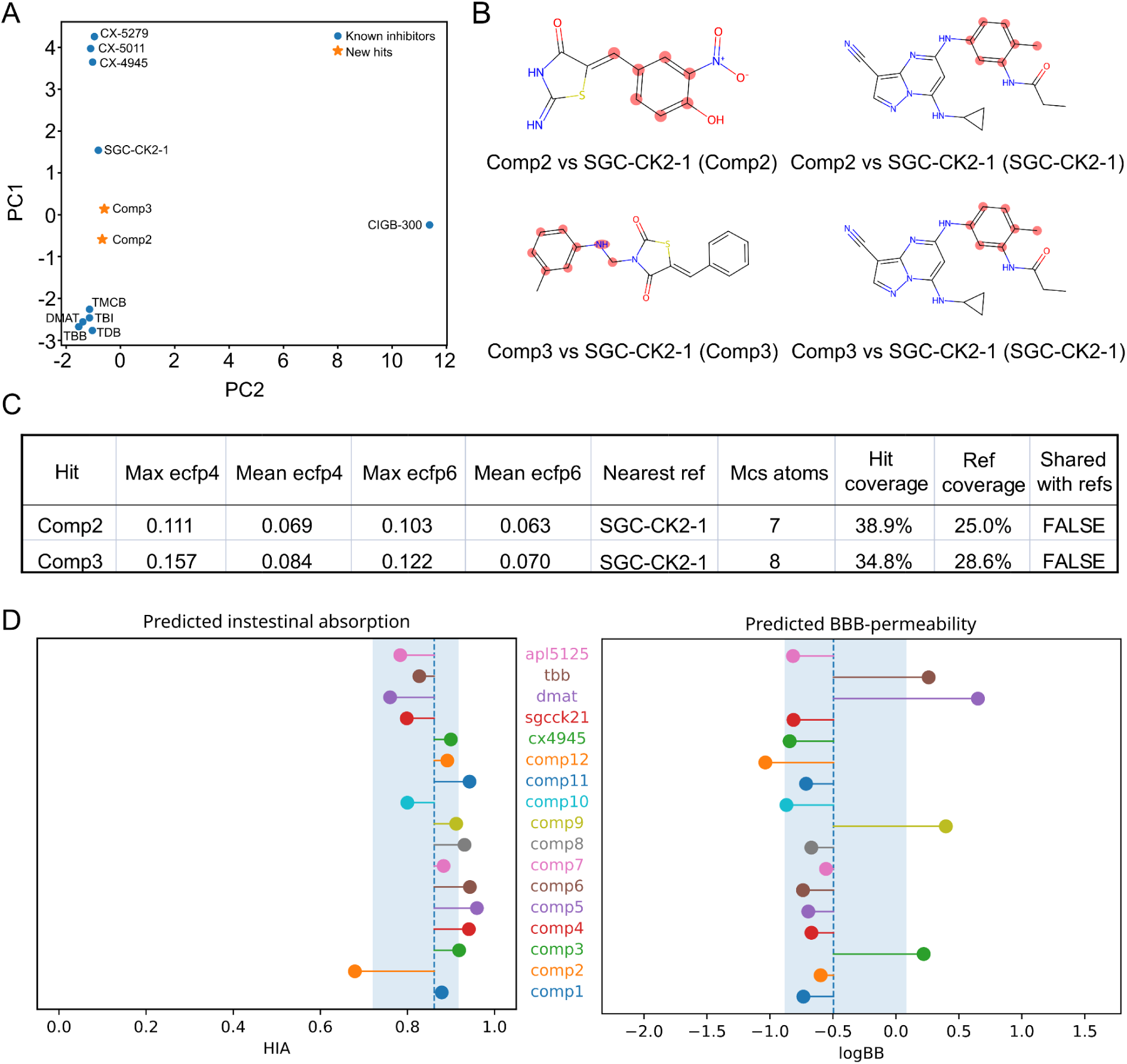
Structural similarity and scaffold novelty of the newly identified CK2 inhibitors. (**A**) Principal component analysis (PCA) of molecular fingerprints (ECFP4) for known CK2 inhibitors (blue circles) and newly identified hits Comp2 and Comp3 (orange stars). (**B**) Maximum common substructure (MCS) comparisons between Comp2 or Comp3 and the most similar known inhibitor (SGC-CK2-1). Shared atoms identified by RDKit FindMCS are highlighted in red. (**C**) Summary of computed molecular similarity metrics for Comp2 and Comp3. Maximum and mean Tanimoto coefficients were calculated based on both ECFP4 and ECFP6 fingerprints. The nearest known inhibitor, number of atoms in the MCS, molecular coverage in each compound, and scaffold overlap status are listed. (**D**) Predicted intestinal absorption and predicted blood-brain barrier permeability of existing and novel CK2 inhibitors. The background data comes from all small-molecule drugs present in the anatomical therapeutic chemical (ATC) classification system (Menestrina et al. 2025). The blue color represents the interquartile range and the dashed line represents the median. Compound 3 is predicted to be both absorbable in the human intestine (HIA) and blood-brain barrier (BBB) permeable.

The maximum common substructure (MCS) analysis further demonstrated limited substructure overlap between the new hits and SGC-CK2-1, with shared substructures consisting of only 7-8 atoms (corresponding to 34-39% molecular coverage in the hits; **Fig. 5B,C**). Similarly, Murcko scaffold comparison, which compares rings and ring-linker structures within small-molecules, confirmed that neither Comp2 nor Comp3 share any scaffold with the reference CK2 inhibitors, indicating that both compounds represent novel chemotypes, distinct from previously established CK2 inhibitor classes (**Figure 5C**). In addition, we performed further chemical analyses of all tested compounds; these analyses show that all tested compounds are drug-like in terms of their physicochemical properties (**Figure S12**), and Comp3 shows structural signals for superior brain bioavailability, over any of the existing CK2 kinase inhibitors (**Figure 5D**).

Together, these findings indicate that ML-predicted compounds 2 and 3 are fully novel CK2 inhibitors, with a distinct scaffold over existing CK2 kinases. While Comp 2 might not be orally bioavailable, the high intestinal absorption and blood-brain penetrance could make Comp3 a potent inhibitor for brain tumours, where CK2 activity sustains cancer cell survival or tumour invasiveness (Pucko and Ostrowski 2022).

## Discussion

Machine intelligence is quickly becoming a core engine in drug discovery, enabling efficient lead identification and speeding-up the whole discovery pipeline (Schneider et al. 2020). Despite the wide adoption, many open problems remain, such as balanced safety and efficacy modelling and prediction. Here, we systematically explored the off-target landscape of human cells using the largest available resources, and showed that molecules with higher affinity towards CDK family off-targets cause more drastic adverse toxicities or failures in clinical trials. By using this off-target knowledge, we designed an ML-powered drug discovery pipeline, which circumvents the key toxicity-inducing targets, while retaining high affinity towards the intended target, CK2. In doing so, we focused both on safety and efficacy aspects of ML-powered drug discovery.

Focusing on molecular safety, we showed that approved pharmaceuticals with high bioactivity towards CDK1/2/7/9 are associated with more severe adverse events. Congruently, drug candidates with activity towards CDK1/2/7/9 have an increased likelihood of being eliminated during the clinical trials. These analyses do not only strengthen our current knowledge of the roles and cellular dependencies of different CDK family members, but they also provide means to perform a more broad-scale and nuanced off-target landscape mapping, covering a wider range of drug targets. We argue that the currently existing drug discovery pipelines fail to appreciate the full spectrum of the anti-target landscape, and broader off-target mapping efforts will form a cornerstone for efficient AI/ML driven drug discovery in the future, enabling more balanced safety/efficacy predictions.

To advance drug efficacy prediction, we coupled a data-centric ML strategy (curated IC_50_ labels from multiple databases, and applicability-domain filtering via an error prediction model) with a structure-specific prioritization to enrich CK2 binders. Using this hybrid pipeline, we attempted to minimize the ML model error with an adjacent error prediction model. To our knowledge, such an error modelling has not been previously applied in drug design, where it was able to increase ML model performance (**Figure S6**), and to deliver a high biochemical hit rate as validated using orthogonal assays. Importantly, the simple ML models, such as SVM, together with the error modelling, seemed superior even to more advanced models such as ANN. This advantage can be largely attributed to the systematic optimization of the dataset prior to model training. Specifically, we performed a multi-dimensional investigation of how different data characteristics influence model performance and selected the most informative feature representations for subsequent training. Such data-centric optimization is rarely emphasized in earlier studies, where substantial effort has often been devoted to increasing model or algorithmic complexity, while the quality and suitability of the input data itself were largely overlooked. Our findings highlight that careful feature engineering and dataset refinement can be at least as critical as model sophistication in achieving robust and generalizable predictive performance.

CK2 protein has multiple suitable binding interfaces for inhibitor design. Here, based on the error modelling with SVM, we were able to predict molecules which either bridged the ATP hinge and the αD pocket (bitopic binding mode) or occupied the αD pocket alone (allosteric inhibition). This is especially reassuring as targeting allosteric regulation sites, which have less sequence and structural conservation across kinases, can help to circumvent major off-target effects or even extend therapeutic target space, compared to active site targeting (Mingione et al. 2023; Fidan et al. 2025). Consistent with previous findings (Bulanova et al. 2024), the newly discovered CK2 inhibitors showed synergistic effects with PARP inhibitors, while being more specific than previous inhibitors as shown using phosphoproteomics and broad kinase-panel profiling.

Study limitations: While this study leverages large-scale open-source bioactivity data, such data remains biased toward well-studied proteins, which may limit applicability of such modelling for other targets. In addition, the models are primarily based on in vitro biochemical measurements and therefore do not fully capture context-dependent cellular effects. Finally, we did not perform experimental structural analysis to confirm small-molecule docking poses, rather we used high-throughput docking as one extra filtering step for our molecular discovery framework. Whereas future research will keep filling the gaps in data availability, laboratories equipped to perform structural protein biology will likely explore the binding of our molecules to CK2.

Taken together, our study delivers three key advances. First, it demonstrates that systematic off-target mapping can be applied prospectively to guide inhibitor design toward safer regions of chemical space. Second, we show that exceptionally high hit rates can be achieved using relatively simple, data-centric machine-learning models, without reliance on deep learning architectures. Third, we identify scaffold-novel and selective CK2 inhibitors for a kinase that currently lacks an FDA-approved inhibitor, and perform a deep experimental validation using orthogonal approaches, including a 216-kinase profiling panel and phosphoproteomics, demonstrating selective suppression of CK2 signaling while avoiding toxicity-inducing CDK family kinases.

## Methods

### Pan-cancer cell line analyses of CK2

Transcriptomic data (log2[TPM+1]) across 33 cancer types were obtained from The Cancer Genome Atlas (TCGA) via the UCSC Xena platform. Cancer type abbreviations and sample details are summarized in **Table S1**. Differential expression analyses of CSNK2A1 and CSNK2A2 between tumor and adjacent normal tissues were performed using Wilcoxon rank-sum tests with false discovery rate (FDR) correction via the Benjamini–Hochberg method, and the results were visualized as boxplots.

For functional dependency assessment, CRISPR-Cas9 gene knockout dependency (CERES) scores and RNA-seq expression data for 1,103 cancer cell lines were retrieved from the DepMap portal (24Q4 Public release). Scatter plots visualizing baseline gene expression versus dependency scores were constructed using ggplot2, with the five most CK2-dependent cell lines (lowest CERES scores) highlighted and annotated via ggrepel. All analyses were performed in R (version 4.4.2), employing packages including ggpubr, ggplot2, survival, forestplot, fmsb, and ggrepel.

### Data collection and preparation

To construct benchmark datasets for building regression models that predict quantitative bioactivities of compounds against casein kinase 2, we collected the drug structures and a wide variety of compound-target bioactivities, including IC_50_, K_d_, K_i_ values from the ChEBML, BindingDB and DrugTargetCommon. Then, we extracted their canonical SMILES using “rdkit.Chem package” in Python and removed the duplicates in each bioactivity type. Finally, 1282 compounds with IC_50_ values, 361 compounds K_i_ and 119 compounds K_d_ values were left.

### Generation of compounds features

To construct an effective regression model, we generated and compared different chemical fingerprints for the compounds. Here, five commonly used fingerprints, including standard fingerprints (1024 bits), estate fingerprints (79 bits), graph fingerprints (1024 bits), MACCS fingerprints (166 bits), and substructure fingerprints (307 bits) were generated using the “rcdk” package in R. The impact of various fingerprints on model prediction accuracy were investigated to get the best input features for the regression model.

### Chemical diversity of the datasets

It is widely recognized that the chemical diversity of a target activity dataset is a crucial factor for building a robust and comprehensive model for predicting compound-target binding affinities. The molecular descriptors including molecular weight (MW), octanol-water partition coefficient (ALogP), topological surface area (TPSA) were calculated using “rdkit.chem.descriptors” package in Python.

### Construction of bioactivity prediction models (BPM)

To predict the bioactivity value of various compounds against CK2, we used the compound fingerprints (standard fingerprint) as the input features in five machine learning algorithms, including random forest (RF), artificial neural network (ANN), support vector machine (SVM), extreme gradient boosting (XGBoost) and gradient boosted decision tree (GBDT), accessible via the scikit python implementation (http://www.scikit-learn.org/). Then, we used the following metrics to score the accuracy of quantitative bioactivity value predictions:

1. Pearson correlation coefficient between the predicted and actual values, which quantifies the linear relationship between the activity predictions.
2. Spearman’s rank correlation coefficient between the predicted and actual values, which quantifies the ability to rank drug pairs in correct order.
3. Normalized Root Mean Square Error (NRMSE): a standardized version of root mean square error (RMSE):

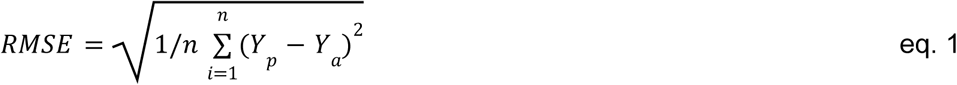

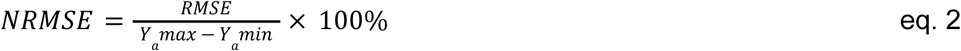

Here, Y_p_ is the predicted bioactivity value and Y_a_ is the measured bioactivity value.

### Construction of error prediction models (EPM)

Model’s application domain refers to the chemical space covered by the training data. To ensure reliable bioactivity predictions, only the compounds located in the application domain of the bioactivity prediction model (BPM) should be used. To evaluate the model application domain, we used an error prediction model (EPM). In this process, the test set was further divided into a calibration set and a new test set using 1:1 ratio. The EPM was trained using the calibration set, and the features of EPM were the same type fingerprints (1024-bit standard fingerprint) as the BPM. The labels of EPM were shown as Equation 3, where Y_p_ is predicted bioactivity value from the BPM, and Y_a_ is actual bioactivity value of these compounds in the calibration set. The machine learning method and parameters of EPM were the same as in the BPM.

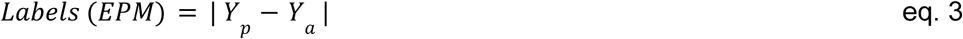

Then, the labels of EPM in the calibration set were defined as the set L, where the values were sorted in ascending order.

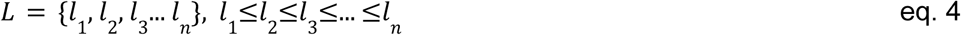

When doing prediction in the new test set, the BPM and EPM were firstly used to get all predicted bioactivity values and error values. Then we could define a confidence level, ɑ. The ɑ-quantile of the L set (Q(ɑ),Equation 5 and 6) in the calibration set would be used as a cutoff to decide which compounds would be left by EPM. The compounds whose predicted error values were less than Q(ɑ) in the new test set would be kept and regarded as reliable predictions.

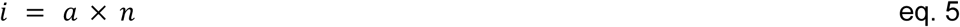

Q(ɑ) was then computed as *l*_*i*_ + *l*_*i*+1_ (if i was an integer), or otherwise as *l_i_* taking the smallest integer greater than or equal to i (ceiling function of i). Here, n is the total number of compounds in the calibration set, eq. 5 is the index corresponding to the α, and li and li+1 are the i-th and (i+1)-th elements in the sorted label set L.

For instance, assuming five calibration samples with ascending error values L={0.1,0.2,0.4,0.6,0.8} and a confidence level α=0.6, the index is i=α×n=3.0. Since i is an integer, the quantile is calculated as Q(0.6)=(l3+l4)/2=(0.4+0.6)/2=0.5. In the subsequent prediction step, if the new test set contains five compounds with predicted error values {0.15,0.25,0.38,0.52,0.70}, those with predicted errors less than Q(α)(e.g., < 0.5) are considered reliable predictions and retained by the EPM, whereas those exceeding this cutoff are excluded.

### Combined BPM and EPM for virtual screening

To virtually screen for compounds with strong affinity towards CK2, we chose a commercial compound library from TargetMol, which contains up to 12 million compounds, including different small molecular types such as FDA approved drugs, novel chemicals, natural products and endogenous metabolites. We generated the standard fingerprint of these compounds as the input features for the IC_50_ bioactivity and error predictions using the BPM and EPM models. After that, a specific Q(ɑ) value was S to select the most accurate predicted compounds and an IC_50_ bioactivity value was used as the threshold to find the compounds with promising affinity to act as the inhibitors of CK2. Finally, Lipinski’s Rule of 5 was applied to select the drug-like compounds.

### The effect of bioactivity value types on regression model accuracy

Various bioactivity types, including IC_50_, K_d_, K_i_, and their combinations, could be used to train regression models. The different bioactivity types used as input will affect the accuracy of the regression models (Cichońska et al. 2021). To find the most suitable bioactivity value types leading to high prediction accuracy, we used each bioactivity value type and their combinations as input to train the regression models. For the combination, when a compound has multiple bioactivity types, the most confident bioactivity type was selected according to the confidence ranking of the various bioactivity categories (Wang et al. 2020). As the number of compounds with K_d_ value was relatively small, we did not use it as the sole bioactivity type to train the models, but rather combined it with other bioactivity types. In addition, apart from the simple combination of different bioactivity types, we also calculated the interaction score (IS) that summarizes IC_50_, K_d_, K_i_ into a single value (represented as IS_biochemical and IS_cell based) (see Tanoli et al. 2020 for interaction score calculations). Compared to other activity types, the interaction score also considers the protein target family and assay format, which could provide more comprehensive information for the compound-target interactions.

After training the 5 regression models with different bioactivity types, we observed that IC_50_ alone or in combination with other bioactivity data types, resulted in highly accurate predictions, especially with K_d_, even if the input dataset size was reduced (**Figure S2**). To further investigate whether the IC_50_ alone or in combination contributed most to the high prediction accuracy, we eliminated dataset-size effects by randomly sub-sampling 1,000 compounds from each bioactivity-type dataset. Then, the datasets were split into a training dataset and a test set with a proportion of 3:1. The models were re-trained in the training dataset and the model prediction accuracy was scored in the test dataset. We observed that the IC_50_ bioactivities alone led to the highest prediction accuracy (**Figure S3**). Thus, we finally selected IC_50_ as the bioactivity value for the model training and prediction.

### The effect of various chemical fingerprints on regression model accuracy

Molecular fingerprints are widely used to describe the structural characteristics of molecules, particularly in relation to their biological activities (Seo et al. 2020). The chemical fingerprints provide an abstract representation of specific structural features. Generated directly from chemical structures, molecular fingerprints can be converted into 2D fragments, enabling an effective representation of chemical fragments in ML models (Zhao et al. 2022). In this study, five commonly used fingerprints with various lengths were compared to characterize the molecules and their impacts on the regression model accuracy. The regression models were trained with different chemical molecule fingerprints as the input feature, and the IC_50_ values as the bioactivity type. After that, the regression model accuracy was scored in the test dataset (**Figure S4**). We observed that most of the chemical fingerprints performed similarly well, but there was a notable decrease on the model prediction accuracy when using the estate fingerprint. As the length of estate fingerprint has only 79 bits, it may be too short to effectively characterize molecules. It has been shown that a limited number of features leads to a significant loss of information, resulting in poor predictions (Xu et al. 2012). Through comparison, the standard fingerprint led to the highest prediction accuracy in most of the regression models, and therefore it was selected as the optimal chemical fingerprint to represent the molecules.

### The effect of molecular similarity on regression model accuracy

To explore the effects of training data size and molecular space diversity that contribute to the regression model accuracy, we calculated the Tanimoto similarity using the optimal fingerprint selected above. Structurally similar compounds were removed based on various thresholds (0.4-1.0). The datasets with different data size and compound similarity were generated and the selected regression model was trained using these datasets. First, for each specific compound in the training set, we calculate its similarity with every compound in the test set. If the maximum similarity is greater than a given threshold, this compound is removed; otherwise, it will be kept in the training set. We tried different thresholds and also performed random removal as shown in **Figure S5**. The SVM method and optimal parameters in grid search were used to establish the models. As the size of the training set decreased in the random removal, the SVM model performance decreased in the test set. This is expected due to the smaller training dataset. The removal by Tanimoto similarity displayed a lower prediction accuracy when compared with random removal, since the training set structural diversity also decreased. These results show that both molecular structural diversity and training sample size are important in this prediction task, and suggest that the model training should be based on as much and as diverse as possible to maximize the model performance in the test set.

### The performance of error prediction model

To evaluate the application domain of bioactivity prediction model (BPM), we set up an error prediction model (EPM) (Sheridan 2013) using SVM method and the same parameters with BPM. The predicted labels of EPM were the errors (Equation 3) from BPM, and the input features of EPM were the standard fingerprints. In this process, the previous test set was divided into a calibration set and a new test set according to the ratio 1:1. Then the calibration set was used to train the EPM. When setting a specific confidence level (ɑ-value), the samples with large predicted error values in the new test set would be removed and we only do bioactivity prediction of left samples. The result is shown in **Figure S6A-D**. To obtain robust results, the above process was repeated 60 times in every confidence level. When using the EPM method, the Spearman coefficient (**Figure S6A**), Pearson coefficient (**Figure S6B**) and RMSE (**Figure S6D**) could achieve significantly superior results. The NRMSE could also decrease when ɑ-value=0.1, 0.2 or 0.3 (**Figure S6C**). However, when ɑ-value was less than 0.5, this metric increased. This was because when only a small part of compounds were left in the new test set, the limited sample size would make the NRMSE have less denominator values (Ya_max_-Ya_min_) in Equation 2, which is not related to model performance. To sum up, NRMSE is a kind of metric related to sample size and EPM could further improve BPM performance by only doing prediction on compounds located in the model application domain.

### High-throughput site-specific molecular docking

Two catalytic isoforms of CK2, designated CK2α (encoded by the CSNK2A1 gene) and CK2α’ (encoded by the CSNK2A2 gene) conjugated with their studied ligands were downloaded from PDB database with high resolutions (CK2α: 5CU4; CK2α’: 8QBU). Using these protein structures, the solvent and other organics were removed from the downloaded proteins, followed by adding the missing hydrogens using AutoDock Tools 1.5.6 software (Eberhardt et al. 2021), and the results were exported in PDBQT format. All small molecule ligands to be docked were converted to PDBQT format and docked using the Vina software. In this process, the ligands were set as flexible structures while the proteins were considered rigid.

The binding energy of each molecular ligand was calculated using the Lamarckian genetic algorithm (Morris et al. 1998), and compounds exhibiting affinity values lower than those of the original inhibitors and effectively binding to the α-D site were screened as potential CK2 inhibitors based on the docking model. Subsequently, the optimal conformations of the selected ligand-protein complexes with the highest binding affinities were visualized using PyMOL.

### Gene dependency analysis using DepMap

The gene dependency analytics was performed from the DepMap release 25Q2. To assess the off-target landscape, gene dependency probabilities for each gene were averaged over all cell models and the mean values were plotted.

### Post-marketing drug severity score

A drug severity (DS) score was defined to estimate the overall post-marketing severity associated with each drug, integrating both the intensity and the fatality of its safety signals while penalizing for concomitant drug use. We used the Individual Case Safety Reports (ICSRs) in the SafetyVista platform (SafetyVista, Chemotargets SL, 2025, https://safety-vista.com/). It includes data from four spontaneous reporting systems (SRS), namely, FAERS (FDA adverse event reporting system. https://open.fda.gov/data/faers/), VAERS (Shimabukuro et al. 2015), VigiBase (Lindquist 2008) and JADER (Pharmaceuticals and Medical Devices Agency. PMDA. https://www.pmda.go.jp/english/index.html). All SRS sources were standardised, de-duplicated and corrected for masking effects, such as those arising from the massive reporting during the COVID-19 pandemic (Montes-Grajales et al. 2023)

For each drug, we first identified all signals of disproportionate reporting (SDRs). An adverse event was considered an SDR if it had 5 or more reports and a value higher than 1 for the lower limit of the 95% confidence interval of proportional reporting ratio (PRR05). Then, for each drug-SDR pair, we calculated the reporting frequency (RF), as well as the proportion of death-related reports (*D*) related to each event. The mean product of these two measures was then divided by the concomitance factor (*C*), defined as the average mean number of drugs reported together with the drug in individual case safety reports. The resulting DS values were normalized between 0 and 1 to allow comparability across drugs.

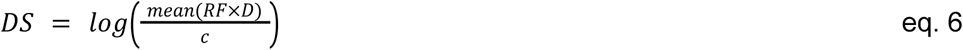

This definition increases the DS score for drugs with frequent and severe (death-associated) safety signals, while decreasing it for drugs whose reports are heavily confounded by concomitant medications.

### Systematic drug-target mapping from multiple databases

Bioactivity data were downloaded as csv files from ChEMBL (version 35) (Zdrazil et al. 2024), BindingDB (1.3.2025 update) (Liu et al. 2025), and Drug Target Commons (accessed March 2025) (Tanoli et al. 2018). A molecule with bioactivity of 1µM or highermore wasis considered as inactive towards a protein target. The following filters were applied on ChEMBL data before the download: Target Organism: Homo sapiens; Molecule Max Phase: Early Phase 1, Phase 1, Phase 2, Phase 3, Approved; Protein Classification L3: Protein Kinase; Target Type: Single protein. After accessing the dataset, we obtained the relevant, CDK-related activities by applying a string search on the Target Name column for “cyclin”, followed by manual filtering of target names. Rows with standard relation of “<” or “>” were removed as the exact activity value is not available. However, if the activity was equal to or greater than 1µM, rows with standard relation “>” were kept due to the molecule being inactive regardless. Rows labeled with “Outside typical range”, “Uncertain”, and “Undetermined”, or with a potential duplicate flag were removed. Nan activity values were filled with 1µM in the presence of a specific comment stating inactivity; otherwise they were removed. The assay confidence scores of the resulting ChEMBL data were 8 or 9 out of 9, indicating high confidence target mapping. For all three databases, only human proteins with activity type IC_50_, EC_50_, K_i_ or K_d_ were considered. For BindingDB and DTC, we required each target based on “UniProt (SwissProt) Recommended Name of Target Chain” and “target_pref_name” to be a CDK, respectively. We mapped BindingDB Monomer IDs to ChEMBL IDs and removed all instances with no matching ChEMBL ID. To avoid redundant data, we mapped all compounds to their parent form based on ChEMBL compound information. Finally, we concatenated all three data frames, computed the geometric mean of duplicate compound-target rows and applied a negative log10 transformation to yield 2,727 unique, CDK-related bioactivity data points. For each CDK, we separated the available bioactivities based on trial phase status; approved drugs in one group and investigational compounds (phases I-III) in another. The groups were further divided by their activity with a threshold of 1µM to form a two-by-two contingency table for testing. The trial phase information of each compound was extracted from ChEMBL (version 35). It should be noted that the trial status of a compound may have changed since.

### Off-target toxicity analysis and Sankey visualization

We selected three representative CK2 inhibitors (CX-4945, DMAT, and TBB) to investigate their off-target interactions with CDKs and related toxicity profiles. Main CDK off-targets of CK2 inhibitors were identified by combining results from ChEMBL, STITCH, and SwissTargetPrediction databases. For each CDK, toxicity associations were retrieved from the Comparative Toxicogenomics Database (CTD) and the top 20 toxicities ranked by Inference Score were selected. The CK2–CDK off-target–toxicity relationships were visualized as a Sankey diagram using the R packages: ggalluvial, ggplot2, and dplyr, which allowed us to map the progression from CK2 inhibitors to CDK off-targets and associated toxicity phenotypes.

### In vitro CK2 kinase assay

In vitro kinase assays were conducted to evaluate the effects of compounds on CK2 activity. Reactions were performed in 384-well plates with a final volume of 10 μL, containing 1.5 ng/μL recombinant CK2 (New England Biolabs, P6010), 50 μM CK2 substrate peptide (Millipore, 12-330), and CK2 assay buffer [40 mM Tris-HCl (pH 7.5), 10 mM MgCl₂, 0.5 mM DTT, and 150 mM NaCl]. DMSO dissolved compounds were added to achieve a final DMSO concentration of 1% (500 nL per well), and the reactions were initiated by adding ATP to a final concentration of 1 μM. Plates were incubated at 37°C for 1.5 hours, after which 10 μL of Kinase-Glo Luminescent Kinase Assay reagent (Promega) was added to quantify residual ATP by luminescence. All tested compounds showed no interference with luciferase activity. IC_50_ values were calculated by fitting dose-response curves using GraphPad Prism.

### 217 kinase activity profiling

Bioactivity profiling of 217 human kinases, encompassing members of the TK, TKL, STE, CMGC, AGC, CAMK, and CK1 families, was performed using the KinaseProfiler platform (ICE Bioscience). For this, compounds were serially diluted from 10 mM DMSO stocks, and transferred into 384-well assay plates using an Echo 655 acoustic dispenser to achieve the desired test concentrations. Kinase selectivity profiling was conducted using both HTRF and ADP-Glo functional assays.

For the HTRF assays, pre-mixed 2× ATP & substrate and 2× kinase/metal ion solutions were sequentially added (2.5 μL each) to the wells containing compounds, followed by incubation at 25 °C for 60 minutes. Subsequently, 5 μL of detection reagent containing XL665 and antibody was added and incubated for another 60 minutes, after which fluorescence signals at 620 nm and 665 nm were recorded using a BMG Labtech microplate reader.

In the ADP-Glo assays, compounds were combined with 2 μL of 2× kinase solution and incubated at 25 °C for 10 minutes, followed by addition of 2 μL of 2× ATP & substrate solution for 60 minutes. ADP-Glo reagent (4 μL) was then added and incubated for 40 minutes, followed by 8 μL of kinase detection reagent for an additional 40-minute incubation prior to luminescence measurement.

Percent inhibition was calculated according to the following equation: % Inhibition = 100% - [(compound - positive control) / (negative control - positive control)] × 100%, where the negative control (1% DMSO) represents 0% inhibition and the positive control (10 μM reference inhibitor) corresponds to 100% inhibition.

### Inhibition test in BRCA cell lines

Eight breast cancer cell lines expressing CK2 were selected based on DepMap transcriptomic profiles (**Table S6**). The 8 lines CAL-120, MCF-7, BT-549, CAL-51, MFM-223, SK-BR-3, MDA-MB-468, and MDA-MB-231, were obtained from the American Type Culture Collection (ATCC, Manassas, VA, USA). CAL-120, BT-549, MDA-MB-231, CAL-51, SK-BR-3 and MFM-223 cells were cultured in RPMI-1640 medium, while MCF-7 and MDA-MB-468 were maintained in DMEM. All media were supplemented with 10% fetal bovine serum (FBS), 5 mM L-glutamine and 1% penicillin-streptomycin, and cells were incubated at 37 °C in a humidified atmosphere containing 5% CO₂. For cytotoxicity assays, cells were seeded in 96-well plates at 5 × 10³ cells per well, allowed to adhere overnight, and then treated with serial dilutions of Compound 2 or Compound 3 (0-400 μM) for 48 hours. Cell viability was determined by MTT assay which is reliable also for chromogenic small molecules (Wang et al. 2024), adding 10 μL of 5 mg/mL MTT solution per well for 4 hours, followed by dissolution in 100 μL DMSO and measurement of absorbance at 570 nm. IC_50_ values were calculated by fitting dose–response curves using GraphPad Prism 10.1.2.

### Drug combination testing

Drugs diluted in dimethyl sulfoxide (DMSO) were dispensed at a 25-nL volume into 384-well black plates (Corning, #3864) using an Echo 550 acoustic liquid handler (Labcyte). CK2 inhibitors (Compound 2 and Compound 3) were tested in combination with PARP inhibitors (rucaparib, olaparib, and niraparib) at seven different concentrations in threefold serial dilutions, covering a concentration range appropriate for each compound. Benzethonium chloride (BzCl, 100 μM) and DMSO (0.1%) were used as positive and negative controls, respectively. SK-BR-3, MDA-MB-231, and MDA-MB-468 breast cancer cells were seeded at a density of 1,000 cells per well and incubated with drug combinations for 72 hours. Cell viability was assessed using the CellTiter-Glo Luminescent Cell Viability Assay (Promega), and luminescence was measured with a Pheras microplate reader. Bliss synergy scores were calculated using SynergyFinder 2.0, with values higher than 8 considered synergistic and negative values interpreted as antagonistic (Ianevski et al. 2020).

### Phosphoproteomics

Drug treated samples (2 hour treatment) were washed with PBS, collected by scraping and snap-freezed. Samples were then lysed in 8 M urea in 50 mM Tris-HCl, pH 8.0, reduced with 10 mM D,L-dithiothreitol, and alkylated with 40 mM iodoacetamide. The samples were digested overnight with sequencing-grade modified Trypsin/Lys-C mix (Promega). After digestion, peptide samples were desalted using a Sep-Pak tC18 96-well plate (Waters) and evaporated to dryness. A small aliquot of the sample was taken for total protein analysis and phosphopeptides were enriched using High Select Fe-NTA Phosphopeptide Enrichment kit (Thermo Scientific). The dried peptides were reconstituted in 0.1% formic acid.

The LC-ESI-MS/MS analyses were performed on a nanoflow HPLC system (Easy-nLC1200, Thermo Scientific) coupled to the Orbitrap Exploris 480 mass spectrometer (Thermo Scientific, Bremen, Germany) equipped with a nano-electrospray ionization source and FAIMS Pro Duo interface (Thermo Scientific, Bremen, Germany). Compensation voltages of -40 V and -60 V were used. Peptides were first loaded on a trapping column and subsequently separated inline on a 15 cm C18 column (75 μm x 15 cm, ReproSil-Pur 3 μm 120 Å C18-AQ, Dr. Maisch HPLC GmbH, Ammerbuch-Entringen, Germany). The mobile phase consisted of water with 0.1% formic acid (solvent A) or acetonitrile/water (80:20 (v/v)) with 0.1% formic acid (solvent B). A 120 min step-gradient was used as follows: from 5% to 21% of solvent B in 62 mins, from 21% to 36% of solvent B in 48 mins, from 36% to 100% in 5 mins, followed by a 5 min wash stage with 100% solvent B.

Samples were analyzed by a data independent acquisition (DIA) LC-MS/MS method. MS data was acquired automatically by using Xcalibur 4.6 software (Thermo Scientific). MS1 spectra were collected in Orbitrap at a resolution of 120000 with a scan range of 395–1005. Normalized AGC target was set to 300% with a maximum injection time of 50 ms. The DIA MS2 scans were acquired with resolution of 30000 over a 400–1000 m/z range and with variable window width. Normalized AGC target was set to 2000% with a maximum injection time of 52 ms.

Data was analysed by Spectronaut software (Biognosys; version 20.2.250922.92449) against *Homo sapiens* database (SwissProt release 2025_03) and Universal Protein Contaminant (Frankenfield et al., 2022) database. DirectDIA approach was used to identify proteins and label-free quantifications were performed with MaxLFQ. Trypsin/P was selected as the enzyme and maximum of 2 missed cleavages were allowed. Carbamidomethylation of cysteine was chosen as fixed modification and protein N-terminal acetylation and oxidation of methionine were selected as variable modifications. For phosphopeptide enriched samples phosphorylation of serine, threonine, and tyrosine were selected as variable modifications. Precursor and protein FDR cutoff were set to 0.01. Quantification was carried out on MS2 level. The data were normalized with the Spectronaut default method, local regression normalization described earlier (Callister et al. 2006).

### Effect of compounds on cellular CK2 activity detected by western blot analysis

SK-BR-3, MDA-MB-468, and MDA-MB-231 breast cancer cells were seeded in 6-well plates at a density of 3 × 10⁵ cells per well and allowed to attach overnight. Cells were then treated with various concentrations of Compound 2 or Compound 3 for 24 hours. Following treatment, cells were washed twice with ice-cold phosphate-buffered saline (PBS) and lysed using RIPA buffer (Thermo Fisher Scientific, USA) supplemented with protease and phosphatase inhibitor cocktails (Sigma-Aldrich, USA). Protein concentrations were determined by the BCA protein assay kit (Pierce, USA) according to the manufacturer’s instructions.

Equal amounts of total protein (30 µg) were resolved by 10% SDS-PAGE and transferred onto polyvinylidene difluoride (PVDF) membranes (Millipore, USA). Membranes were blocked with 5% non-fat dry milk in Tris-buffered saline containing 0.1% Tween-20 (TBST) for 1 hour at room temperature, followed by overnight incubation at 4 °C with primary antibodies against phospho-CK2 substrate (1:1000 dilution, Cell Signaling Technology, #8738) and GAPDH (1:10000 dilution, Abcam) as a loading control. After washing three times with TBST, membranes were incubated with appropriate horseradish peroxidase (HRP)-conjugated secondary antibodies (1:5000, Cell Signaling Technology) for 1 hour at room temperature. Protein bands were visualized using an enhanced chemiluminescence (ECL) detection kit (Thermo Fisher Scientific) and imaged with a ChemiDoc XRS+ system (Bio-Rad, USA). Band intensities were quantified using ImageJ software (NIH), and phospho-CK2 substrate signals were normalized to GAPDH.

### Analysis of structural similarity and scaffold novelty

To evaluate the structural novelty of the newly identified CK2 inhibitors (Comp2 and Comp3) relative to known CK2 inhibitors, a cheminformatics workflow was implemented in Python using RDKit (v2022.09). We collected ten commonly-reported CK2 inhibitors (**Table S5**) as references and all the compounds including our two new hits (Comp2 and Comp3) were compiled and standardized. For each compound, extended-connectivity fingerprints (ECFPs) were calculated using the Morgan algorithm at two different radii (ECFP4, radius = 2; and ECFP6, radius = 3; 2048-bit). Pairwise Tanimoto similarity coefficients were computed between each new hit and all known CK2 inhibitors, and both the maximum and mean similarity scores were recorded. Then, principal component analysis (PCA) was performed on the ECFP4 fingerprints using scikit-learn (v1.4) to visualize the overall distribution of chemical space. The first two principal components (PC1 and PC2) were plotted to generate a two-dimensional map, with known inhibitors shown as blue circles and new hits as orange stars. To assess scaffold novelty, Bemis-Murcko scaffolds were extracted for all compounds, and the scaffolds of Comp2 and Comp3 were compared with those of the reference inhibitors. Compounds that did not share any scaffold with known inhibitors were considered to possess novel structural frameworks.

For each hit, the most similar known CK2 inhibitor (based on the highest ECFP4 similarity) was selected for maximum common substructure (MCS) analysis. MCS identification was performed using the RDKit FindMCS function with “ringMatchesRingOnly” and “completeRingsOnly” constraints. The number of shared atoms and molecular coverage for both compounds were calculated, and the matched atoms were visualized as highlighted fragments on 2D molecular diagrams. All analyses and visualizations were performed using RDKit, pandas (v2.2.2), numpy (v1.26.4), matplotlib (v3.9.2), and scikit-learn (v1.5.1).

## Supporting information

Supplementary material including supplementary figures

Large version of supplementary figure 9

## Data availability

All processed datasets used for machine learning model training and evaluation are included in the GitHub repository (https://github.com/YingHHBio/CK2-ML-screening). The other raw data will be made available upon reasonable request to the corresponding author.

## Code availability

All codes to produce the analyses as shown in this publication are available through GitHub (https://github.com/YingHHBio/CK2-ML-screening).

## Acknowledgements

We want to thank Maria Taskinen and Henri Xhaard for their expert comments during the writing of this manuscript. The authors acknowledge the CSC – IT Center for Science, Finland, for providing the computational resources. Mass spectrometry analysis was performed at the Turku Proteomics Facility, University of Turku and Åbo Akademi University. The facility is supported by Biocenter Finland. Drug testing was carried out at the FIMM High Throughput Biomedicine Unit, hosted by the University of Helsinki and supported by HiLIFE and Biocenter Finland. Funding support: T. Aittokallio: Research Council of Finland (grants 344698, 345803, and 367855); the Cancer Foundation of Finland, the Norwegian Cancer Society (grant 273810), the Sigrid Jusélius Foundation, and iCAN – Digital Precision Cancer Medicine Flagship (iCAN-MULTIDRUG). H. Ying was supported by grants from K. Albin Johansson Stiftelse Foundation (No. 2023–12), the Maud Kuistila Memorial Foundation sr (No. 2024-0147B), the Finnish Cultural Foundation (No. 00251048), Jalmari ja Rauha Ahokkaan Säätiö foundation, Paulon Säätiö foundation and the China Scholarship Council (No. 2020-I78).

## Notes

https://github.com/mitroe/CK2_kinase_inhibitor_prediction/

### Competing Interest Statement

Jordi Mestres is the founder and research director of Chemotargets. Mitro Miihkinen and Tero Aittokallio receive research and salary funding from Mobius biotechnology. The authors declare no other competing interests.

### Summary of Updates

New figure 5 added characterising molecular properties of new compounds. New phosphoproteomic analyses added on Figure 4 and improved all figures altogether. Removed multiple typoes.

## References

Ahlmann-Eltze, Constantin, Wolfgang Huber, and Simon Anders. 2025. ‘Deep-Learning-Based Gene Perturbation Effect Prediction Does Not yet Outperform Simple Linear Baselines’. Nature Methods 22 (8): 1657–61. 10.1038/s41592-025-02772-6.

Attwood, Misty M., Doriano Fabbro, Aleksandr V. Sokolov, Stefan Knapp, and Helgi B. Schiöth. 2021. ‘Trends in Kinase Drug Discovery: Targets, Indications and Inhibitor Design’. Nature Reviews Drug Discovery 20 (11): 839–61. 10.1038/s41573-021-00252-y.

Bilodeau, Stéphanie, Véronique Caron, Jonathan Gagnon, Alexandre Kuftedjian, and André Tremblay. 2017. ‘A CK2–RNF4 Interplay Coordinates Non-Canonical SUMOylation and Degradation of Nuclear Receptor FXR’. Journal of Molecular Cell Biology 9 (3): 195–208. 10.1093/jmcb/mjx009.

Borgo, Christian, Luca Cesaro, Tsuyoshi Hirota, et al. 2021. ‘Comparing the Efficacy and Selectivity of Ck2 Inhibitors. A Phosphoproteomics Approach’. European Journal of Medicinal Chemistry 214 (March): 113217. 10.1016/j.ejmech.2021.113217.

Borgo, Christian, Claudio D’Amore, Stefania Sarno, Mauro Salvi, and Maria Ruzzene. 2021. ‘Protein Kinase CK2: A Potential Therapeutic Target for Diverse Human Diseases’. Signal Transduction and Targeted Therapy 6 (May): 183. 10.1038/s41392-021-00567-7.

Bowes, Joanne, Andrew J. Brown, Jacques Hamon, et al. 2012. ‘Reducing Safety-Related Drug Attrition: The Use of in Vitro Pharmacological Profiling’. Nature Reviews Drug Discovery 11 (12): 909–22. 10.1038/nrd3845.

Brear, Paul, Darby Ball, Katherine Stott, Sheena D’Arcy, and Marko Hyvönen. 2020. ‘Proposed Allosteric Inhibitors Bind to the ATP Site of CK2α’. Journal of Medicinal Chemistry 63 (21): 12786–98. 10.1021/acs.jmedchem.0c01173.

Brennan, Richard J., Stephen Jenkinson, Andrew Brown, et al. 2024. ‘The State of the Art in Secondary Pharmacology and Its Impact on the Safety of New Medicines’. Nature Reviews Drug Discovery 23 (7): 525–45. 10.1038/s41573-024-00942-3.

Bulanova, Daria, Yevhen Akimov, Wojciech Senkowski, et al. 2024. ‘A Synthetic Lethal Dependency on Casein Kinase 2 in Response to Replication-Perturbing Therapeutics in RB1-Deficient Cancer Cells’. Science Advances 10 (21): eadj1564. 10.1126/sciadv.adj1564.

Callister, Stephen J., Richard C. Barry, Joshua N. Adkins, et al. 2006. ‘Normalization Approaches for Removing Systematic Biases Associated with Mass Spectrometry and Label-Free Proteomics’. Journal of Proteome Research 5 (2): 277–86. 10.1021/pr050300l.

Cichońska, Anna, Balaguru Ravikumar, Robert J. Allaway, et al. 2021. ‘Crowdsourced Mapping of Unexplored Target Space of Kinase Inhibitors’. Nature Communications 12 (June): 3307. 10.1038/s41467-021-23165-1.

Eberhardt, Jerome, Diogo Santos-Martins, Andreas F. Tillack, and Stefano Forli. 2021. ‘AutoDock Vina 1.2.0: New Docking Methods, Expanded Force Field, and Python Bindings’. Journal of Chemical Information and Modeling, ahead of print, July 19. World. 10.1021/acs.jcim.1c00203.

Elmore, Andrew R., Aws Sadik, Lavinia Paternoster, Golam M. Khandaker, Tom R. Gaunt, and Gibran Hemani. 2025. ‘Genetic Inference of On-Target and off-Target Side-Effects of Antipsychotic Medications’. PLOS Genetics 21 (7): e1011793. 10.1371/journal.pgen.1011793.

Fidan, Vahap Gazi, Konuralp Ilim, Attila Gursoy, S. Banu Ozkan, and Ozlem Keskin. 2025. ‘Rewiring Enzyme Regulation: Allosteric Drugs and Predictive Tools’. Current Opinion in Structural Biology 95 (December): 103159. 10.1016/j.sbi.2025.103159.

Gandini, Annachiara, Manuela Bartolini, Daniele Tedesco, et al. 2018. ‘Tau-Centric Multitarget Approach for Alzheimer’s Disease: Development of First-in-Class Dual Glycogen Synthase Kinase 3β and Tau-Aggregation Inhibitors’. Journal of Medicinal Chemistry 61 (17): 7640–56. 10.1021/acs.jmedchem.8b00610.

Giri, Anil K., Aleksandr Ianevski, and Tero Aittokallio. 2019. ‘Genome-Wide off-Targets of Drugs: Risks and Opportunities’. Cell Biology and Toxicology 35 (6): 485–87. 10.1007/s10565-019-09491-7.

Goldwaser, Elodie, Catherine Laurent, Nathalie Lagarde, et al. 2022. ‘Machine Learning-Driven Identification of Drugs Inhibiting Cytochrome P450 2C9’. PLOS Computational Biology 18 (1): e1009820. 10.1371/journal.pcbi.1009820.

Han, L. Y., X. H. Ma, H. H. Lin, et al. 2008. ‘A Support Vector Machines Approach for Virtual Screening of Active Compounds of Single and Multiple Mechanisms from Large Libraries at an Improved Hit-Rate and Enrichment Factor’. Journal of Molecular Graphics & Modelling 26 (8): 1276–86. 10.1016/j.jmgm.2007.12.002.

Ianevski, Aleksandr, Anil K. Giri, and Tero Aittokallio. 2020. ‘SynergyFinder 2.0: Visual Analytics of Multi-Drug Combination Synergies’. Nucleic Acids Research 48 (W1): W488–93. 10.1093/nar/gkaa216.

Ianevski, Aleksandr, Kristen Nader, Kyriaki Driva, et al. 2024. ‘Single-Cell Transcriptomes Identify Patient-Tailored Therapies for Selective Co-Inhibition of Cancer Clones’. Nature Communications 15 (1): 8579. 10.1038/s41467-024-52980-5.

Iegre, Jessica, Eleanor L. Atkinson, Paul D. Brear, Bethany M. Cooper, Marko Hyvönen, and David R. Spring. 2021. ‘Chemical Probes Targeting the Kinase CK2: A Journey Outside the Catalytic Box’. Organic & Biomolecular Chemistry 19 (20): 4380–96. 10.1039/d1ob00257k.

Isigkeit, Laura, Tim Hörmann, Espen Schallmayer, et al. 2024. ‘Automated Design of Multi-Target Ligands by Generative Deep Learning’. Nature Communications 15 (1): 7946. 10.1038/s41467-024-52060-8.

Jorissen, Robert N., and Michael K. Gilson. 2005. ‘Virtual Screening of Molecular Databases Using a Support Vector Machine’. Journal of Chemical Information and Modeling 45 (3): 549–61. 10.1021/ci049641u.

Kufareva, Irina, Benoit Bestgen, Paul Brear, et al. 2019. ‘Discovery of Holoenzyme-Disrupting Chemicals as Substrate-Selective CK2 Inhibitors’. Scientific Reports 9 (1): 15893. 10.1038/s41598-019-52141-5.

Liang, Jing-Wei, Ming-Yang Wang, Shan Wang, Shi-Long Li, Wan-Qiu Li, and Fan-Hao Meng. 2020. ‘Identification of Novel CDK2 Inhibitors by a Multistage Virtual Screening Method Based on SVM, Pharmacophore and Docking Model’. Journal of Enzyme Inhibition and Medicinal Chemistry 35 (1): 235–44. 10.1080/14756366.2019.1693702.

Lindquist, Marie. 2008. ‘VigiBase, the WHO Global ICSR Database System: Basic Facts’. Drug Information Journal : DIJ / Drug Information Association 42 (5): 409–19. 10.1177/009286150804200501.

Liu, Changchang, Peter Kutchukian, Nhan D. Nguyen, Mohammed AlQuraishi, and Peter K. Sorger. 2023. ‘A Hybrid Structure-Based Machine Learning Approach for Predicting Kinase Inhibition by Small Molecules’. Journal of Chemical Information and Modeling, ahead of print, August 18. World. 10.1021/acs.jcim.3c00347.

Liu, Tiqing, Linda Hwang, Stephen K. Burley, et al. 2025. ‘BindingDB in 2024: A FAIR Knowledgebase of Protein-Small Molecule Binding Data’. Nucleic Acids Research 53 (D1): D1633–44. 10.1093/nar/gkae1075.

Menestrina, Luca, Raquel Parrondo-Pizarro, Ismael Gómez, Ricard Garcia-Serna, Scott Boyer, and Jordi Mestres. 2025. ‘Refined ADME Profiles for ATC Drug Classes’. Pharmaceutics 17 (3): 308. 10.3390/pharmaceutics17030308.

Mingione, Victoria R., YiTing Paung, Ian R. Outhwaite, and Markus A. Seeliger. 2023. ‘Allosteric Regulation and Inhibition of Protein Kinases’. Biochemical Society Transactions 51 (1): 373–85. 10.1042/BST20220940.

Montes-Grajales, Diana, Ricard Garcia-Serna, and Jordi Mestres. 2023. ‘Impact of the COVID-19 Pandemic on the Spontaneous Reporting and Signal Detection of Adverse Drug Events | Scientific Reports’.

Morris, Garrett M., David S. Goodsell, Robert S. Halliday, et al. 1998. ‘Automated Docking Using a Lamarckian Genetic Algorithm and an Empirical Binding Free Energy Function’. Journal of Computational Chemistry 19 (14): 1639–62. 10.1002/(SICI)1096-987X(19981115)19:14%253C1639::AID-JCC10%253E3.0.CO;2-B.

Pizzi, Marco, Francesco Piazza, Claudio Agostinelli, et al. 2015. ‘Protein Kinase CK2 Is Widely Expressed in Follicular, Burkitt and Diffuse Large B-Cell Lymphomas and Propels Malignant B-Cell Growth’. Oncotarget 6 (9): 6544–52. 10.18632/oncotarget.3446.

Pucko, Emanuela B., and Robert P. Ostrowski. 2022. ‘Inhibiting CK2 among Promising Therapeutic Strategies for Gliomas and Several Other Neoplasms’. Pharmaceutics 14 (2): 331. 10.3390/pharmaceutics14020331.

Rogers, David, and Mathew Hahn. 2010. ‘Extended-Connectivity Fingerprints’. Journal of Chemical Information and Modeling 50 (5): 742–54. 10.1021/ci100050t.

Scaglioni, Pier Paolo, Thomas M. Yung, Lu Fan Cai, et al. 2006. ‘A CK2-Dependent Mechanism for Degradation of the PML Tumor Suppressor’. Cell 126 (2): 269–83. 10.1016/j.cell.2006.05.041.

Schmidt, Friedemann, Richard J. Brennan, Steve Jenkinson, and Jean-Pierre Valentin. 2025. ‘Shaping Secondary Pharmacology Panels of the Future: Evolving Target Selection Criteria for Safety Panels’. Nature Reviews Drug Discovery 24 (6): 482–84. 10.1038/s41573-025-01184-7.

Schneider, Petra, W. Patrick Walters, Alleyn T. Plowright, et al. 2020. ‘Rethinking Drug Design in the Artificial Intelligence Era’. Nature Reviews Drug Discovery 19 (5): 353–64. 10.1038/s41573-019-0050-3.

Schulman, Aron, Juho Rousu, Tero Aittokallio, and Ziaurrehman Tanoli. 2024. Attention-Based Approach to Predict Drug–Target Interactions across Seven Target Superfamilies. 10.1093/bioinformatics/btae496.

Seo, Myungwon, Hyun Kil Shin, Yoochan Myung, Sungbo Hwang, and Kyoung Tai No. 2020. ‘Development of Natural Compound Molecular Fingerprint (NC-MFP) with the Dictionary of Natural Products (DNP) for Natural Product-Based Drug Development’. Journal of Cheminformatics 12 (1): 6. 10.1186/s13321-020-0410-3.

Sheridan, Robert P. 2013. ‘Using Random Forest to Model the Domain Applicability of Another Random Forest Model’. Journal of Chemical Information and Modeling 53 (11): 2837–50. 10.1021/ci400482e.

Shimabukuro, Tom T., Michael Nguyen, David Martin, and Frank DeStefano. 2015. ‘Safety Monitoring in the Vaccine Adverse Event Reporting System (VAERS)’. Vaccine 33 (36): 4398–405. 10.1016/j.vaccine.2015.07.035.

Tanoli, Ziaurrehman, Zaid Alam, Aleksandr Ianevski, Krister Wennerberg, Markus Vähä-Koskela, and Tero Aittokallio. 2020. ‘Interactive Visual Analysis of Drug-Target Interaction Networks Using Drug Target Profiler, with Applications to Precision Medicine and Drug Repurposing’. Briefings in Bioinformatics 21 (1): 211–20. 10.1093/bib/bby119.

Tanoli, ZiaurRehman, Zaid Alam, Markus Vähä-Koskela, et al. 2018. ‘Drug Target Commons 2.0: A Community Platform for Systematic Analysis of Drug–Target Interaction Profiles’. Database 2018 (January). 10.1093/database/bay083.

Tanoli, Ziaurrehman, Adrià Fernández-Torras, Umut Onur Özcan, et al. 2025. ‘Computational Drug Repurposing: Approaches, Evaluation of in Silico Resources and Case Studies’. Nature Reviews. Drug Discovery 24 (7): 521–42. 10.1038/s41573-025-01164-x.

Tanoli, Ziaurrehman, Aron Schulman, and Tero Aittokallio. 2025. ‘Validation Guidelines for Drug-Target Prediction Methods’. Expert Opinion on Drug Discovery 20 (1): 31–45. 10.1080/17460441.2024.2430955.

Theisen, Ryan, Tianduanyi Wang, Balaguru Ravikumar, Rayees Rahman, and Anna Cichońska. 2024. ‘Leveraging Multiple Data Types for Improved Compound-Kinase Bioactivity Prediction’. Nature Communications 15 (1): 7596. 10.1038/s41467-024-52055-5.

Tibo, Alessandro, Jiazhen He, Jon Paul Janet, Eva Nittinger, and Ola Engkvist. 2024. ‘Exhaustive Local Chemical Space Exploration Using a Transformer Model’. Nature Communications 15 (1): 7315. 10.1038/s41467-024-51672-4.

Trott, Oleg, and Arthur J. Olson. 2010. ‘AutoDock Vina: Improving the Speed and Accuracy of Docking with a New Scoring Function, Efficient Optimization and Multithreading’. Journal of Computational Chemistry 31 (2): 455–61. 10.1002/jcc.21334.

Wang, Sheng, Sebastian O. Klein, Sylvia Urban, et al. 2024. ‘Structure-Guided Design of a Selective Inhibitor of the Methyltransferase KMT9 with Cellular Activity’. Nature Communications 15 (1): 43. 10.1038/s41467-023-44243-6.

Wang, Tianduanyi, Prson Gautam, Juho Rousu, and Tero Aittokallio. 2020. ‘Systematic Mapping of Cancer Cell Target Dependencies Using High-Throughput Drug Screening in Triple-Negative Breast Cancer’. Computational and Structural Biotechnology Journal 18 (January): 3819–32. 10.1016/j.csbj.2020.11.001.

Xu, Congying, Feixiong Cheng, Lei Chen, et al. 2012. ‘In Silico Prediction of Chemical Ames Mutagenicity’. Journal of Chemical Information and Modeling 52 (11): 2840–47. 10.1021/ci300400a.

Zahoránszky-Kőhalmi, Gergely, Cristian G. Bologa, and Tudor I. Oprea. 2016. ‘Impact of Similarity Threshold on the Topology of Molecular Similarity Networks and Clustering Outcomes’. Journal of Cheminformatics 8 (March): 16. 10.1186/s13321-016-0127-5.

Zdrazil, Barbara, Eloy Felix, Fiona Hunter, et al. 2024. ‘The ChEMBL Database in 2023: A Drug Discovery Platform Spanning Multiple Bioactivity Data Types and Time Periods’. Nucleic Acids Research 52 (D1): D1180–92. 10.1093/nar/gkad1004.

Zhang, Kai-ping, Bao-feng Yang, and Bao-xin Li. 2014. ‘Translational Toxicology and Rescue Strategies of the hERG Channel Dysfunction: Biochemical and Molecular Mechanistic Aspects’. Acta Pharmacologica Sinica 35 (12): 1473–84. 10.1038/aps.2014.101.

Zhao, Xia, Yuhao Sun, Ruiqiu Zhang, et al. 2022. ‘Machine Learning Modeling and Insights into the Structural Characteristics of Drug-Induced Neurotoxicity’. Journal of Chemical Information and Modeling 62 (23): 6035–45. 10.1021/acs.jcim.2c01131.

