## Supplementary material including supplementary figures for "Off-target mapping enhances selectivity of machine learning-predicted CK2 inhibitors"

**<u>Descriptions of supplementary tables:</u>**

**Supplementary Table S1.** The list of 33 cancer types and their abbreviations in the pan-cancer analysis from The Cancer Genome Atlas (TCGA) used in this study.

**Supplementary Table S2.** The reported binding affinities ($IC_{50}/K_d$) of CX-4945 toward CDK family kinases retrieved from public kinase profiling databases (ChEMBL, BindingDB).

**Supplementary Table S3.** Kinase profiling results of compound 2 against a 217-kinase panel. Note:A negative inhibition rate indicates that the compound has no inhibitory effect on the kinase.

**Supplementary Table S4.** Kinase profiling results of compound 3 against a 217-kinase panel. Note:A negative inhibition rate indicates that the compound has no inhibitory effect on the kinase.

**Supplementary Table S5.** Existing CK2 inhibitors.

**Supplementary Table S6.** DepMap (Public 24Q4) Chronos CRISPR dependency scores and expression levels of CK2 catalytic subunits (CSNK2A1/CSNK2A2) in selected breast cancer cell lines.

### Supplementary figures 1-12:

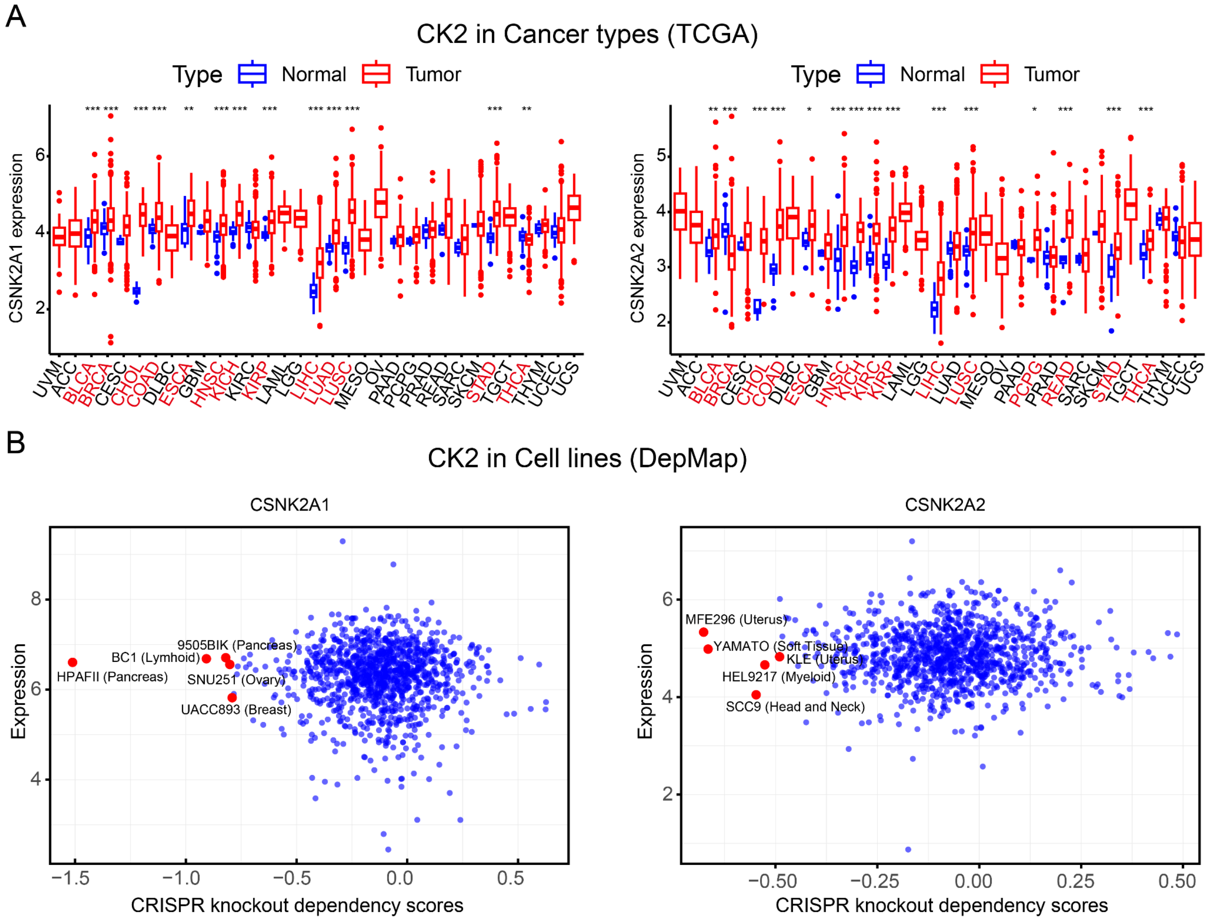


**Supplementary Figure 1**. **Cancer genetics of CK2. (A)** Pan-cancer expression of CK2 genes (CSNK2A1 and CSNK2A2). Differential expression analysis of CSNK2A1 and CSNK2A2 across 33 cancer types in TCGA. Box plots compare tumor (red) and adjacent normal tissues (blue). Cancer types with statistically significant differential expression between tumor and normal tissue (Wilcoxon test, FDR $p < 0.05$) are highlighted in red. **(B)** Cancer cell line dependency on CK2 genes (CSNK2A1 and CSNK2A2). Scatter plots of showing baseline gene expression levels (log2[TPM + 1]) versus CRISPR knockout dependency scores for CSNK2A1 and CSNK2A2 across 1,103 cancer cell lines from different tissue lineages (DepMap, https://depmap.org/portal/). Five cell lines with strong CK2 dependency (more negative CRISPR scores) are highlighted in red and annotated with their cell line name and lineage. This plot illustrates how baseline expression levels relate to cell growth dependency upon gene knockout.

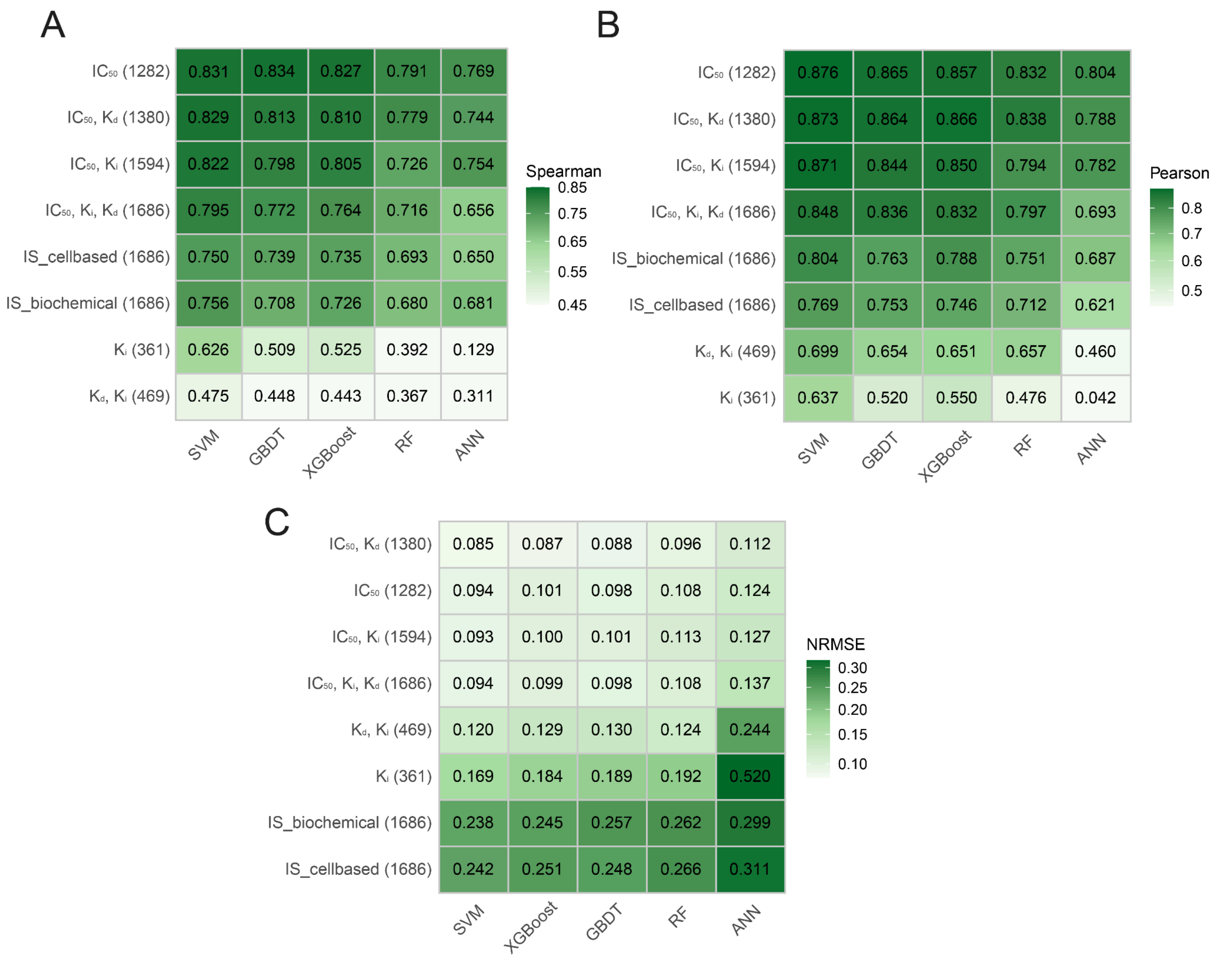


**Supplementary Figure 2. ML model prediction accuracy in the test dataset when trained using various bioactivity types.** The total number of compounds in the datasets is written in parentheses. **(A)** Spearman correlation **(B)** Pearson correlation **(C)** Normalized Root Mean Square Error (NRMSE). The datasets were split into a training dataset and a test set with a proportion of 3:1. IS_biochemical and IS_cellbased are normalized interaction scores ranging from 0 to 1, where 1 indicates the highest potency and 0 indicates no detectable potency.

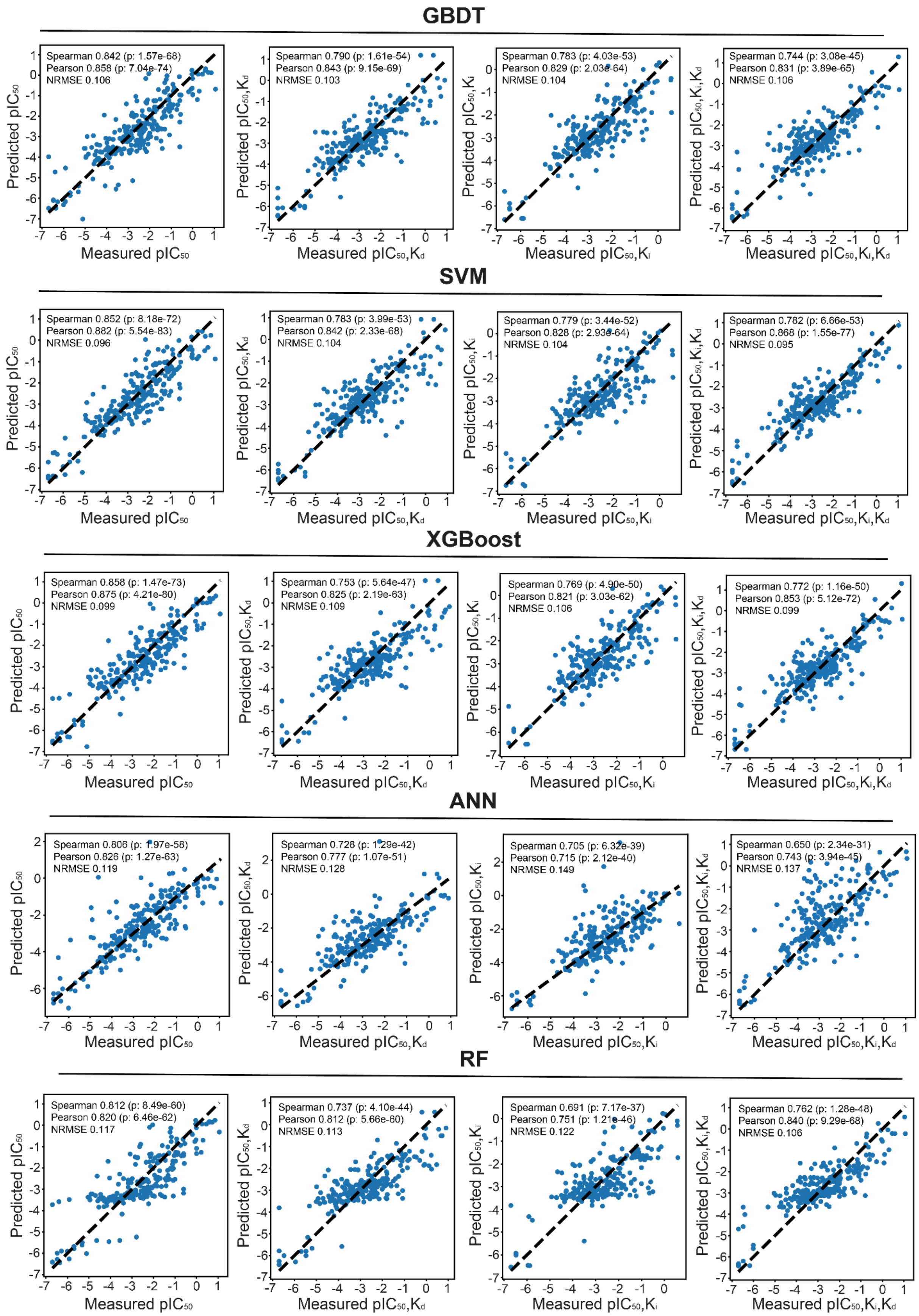


**Supplementary Figure 3**. Scatter plot between predicted and measured bioactivity values in test set (250 compounds) based on the regression models trained with selected bioactivity value types. 1000 compounds in each dataset were randomly selected from the full datasets with different bioactivity values ($IC_{50}$ only; combined bioactivity values of $IC_{50}$, $K_d$; combined bioactivity values of $IC_{50}$, $K_i$; combined bioactivity values of $IC_{50}$, $K_i$, $K_d$).

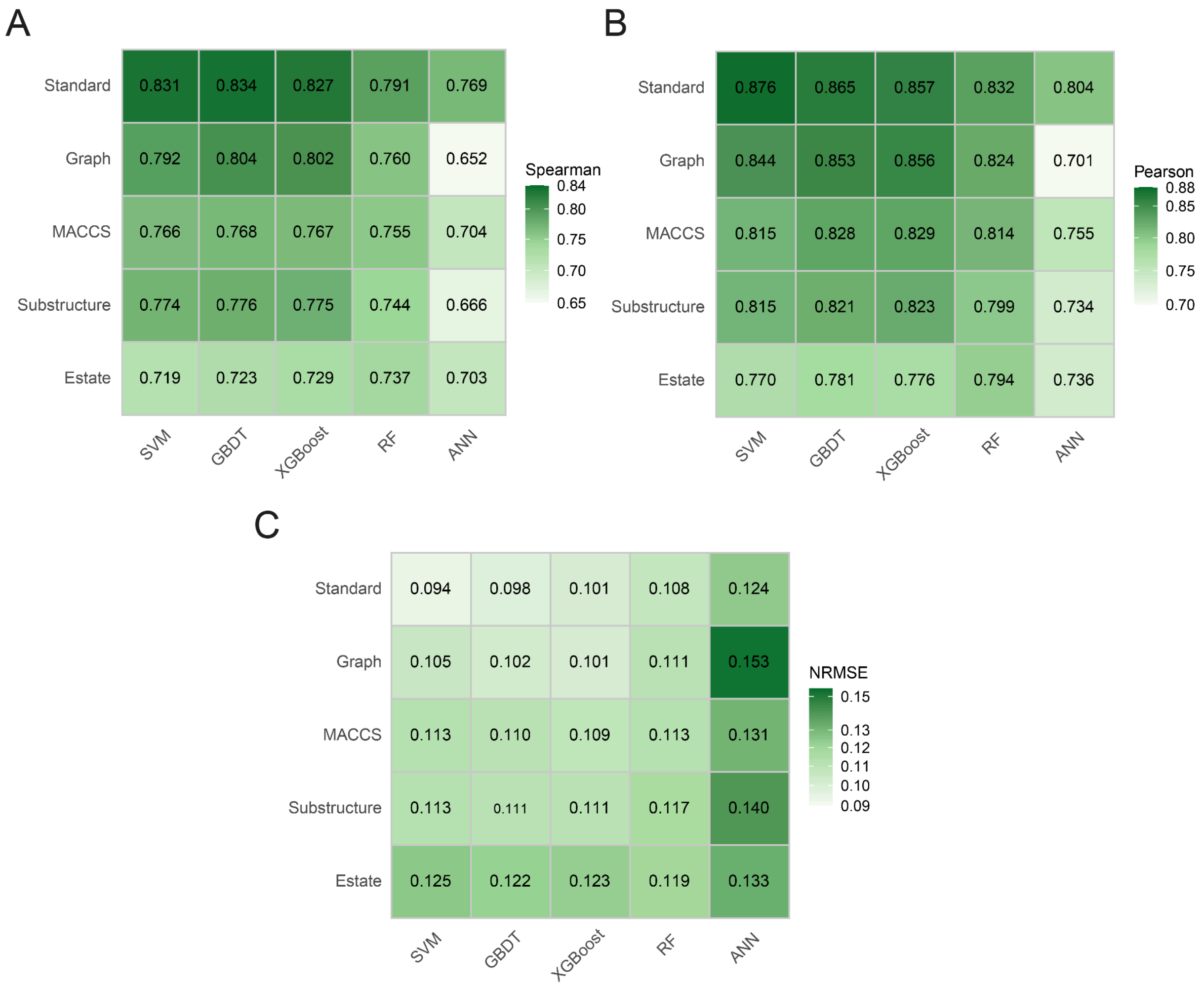


**Supplementary Figure 4**. **The ML model prediction accuracy in the test dataset when using various molecular fingerprints as input features to train the regression models. (A)** Spearman correlation **(B)** Pearson correlation **(C)** Normalized Root Mean Square Error (NRMSE). The datasets were split into a training dataset and a test set with a proportion of 3:1.

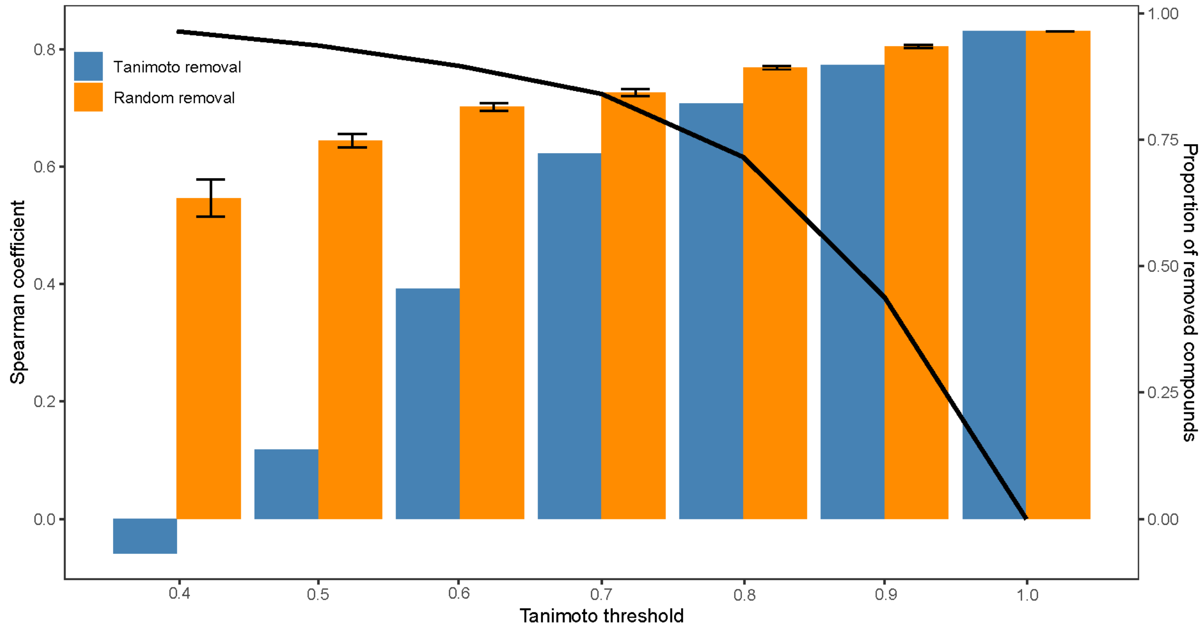


**Supplementary Figure 5**. The SVM model performance as a function of training data size and molecular similarity. In the random removal, the proportion of removed compounds was kept the same as in removal by the Tanimoto similarity. Every random removal was repeated 20 times. The curve show the proportion of removed compounds in the training set (right y-axis).

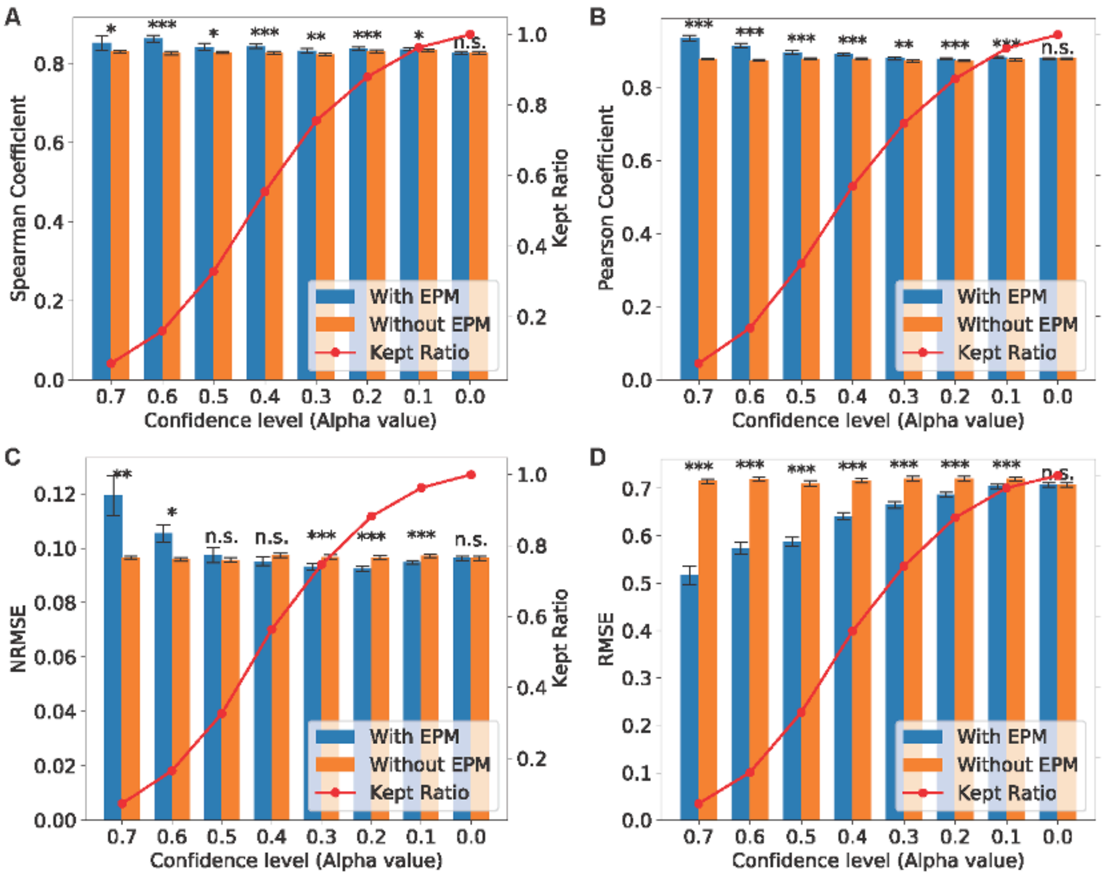

**Supplementary Figure 6**. **The error prediction model performance.** Model performance was assessed using the new set after splitting the test set into a calibration set and a new test set. Dataset split and error model construction were repeated 60 times at every confidence level. The Wilcoxon test was used to calculate p-values. *p<0.05, **p<0.01, ***p<0.001. (**A–B**) Spearman and Pearson correlation coefficients across confidence levels. (**C–D**) NRMSE and RMSE of synergy prediction across confidence levels. The x-axis indicates the confidence level (α), and the left y-axis shows the corresponding performance metric. Blue bars represent results with the error prediction model (EPM), and orange bars represent results without EPM. The red line (right y-axis) denotes the kept ratio, defined as the proportion of samples retained after confidence-based filtering.

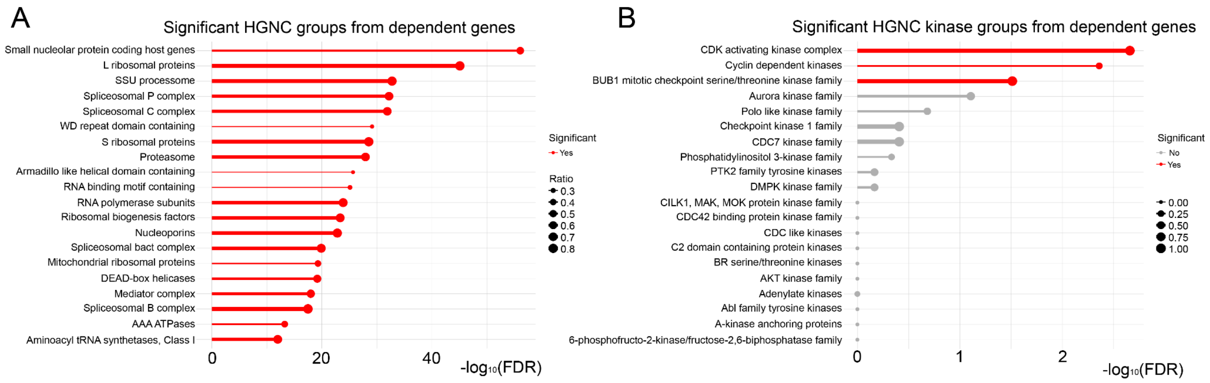


**Supplementary Figure 7**. **Overrepresented protein groups among genes most dependent for cell growth and survival.** (**A**) Top 20 most significant HUGO Gene Nomenclature Committee (HGNC) groups and (**B**) Top 20 most significant kinase-specific groups overrepresented in the dependent gene list. The thickness of the segments corresponds to the ratio of the number of genes found in our list vs the total number of genes present in the HGNC group. Out of 102 kinase-specific HGNC groups, only 3 are significant and they all correspond to CDK-related functions.

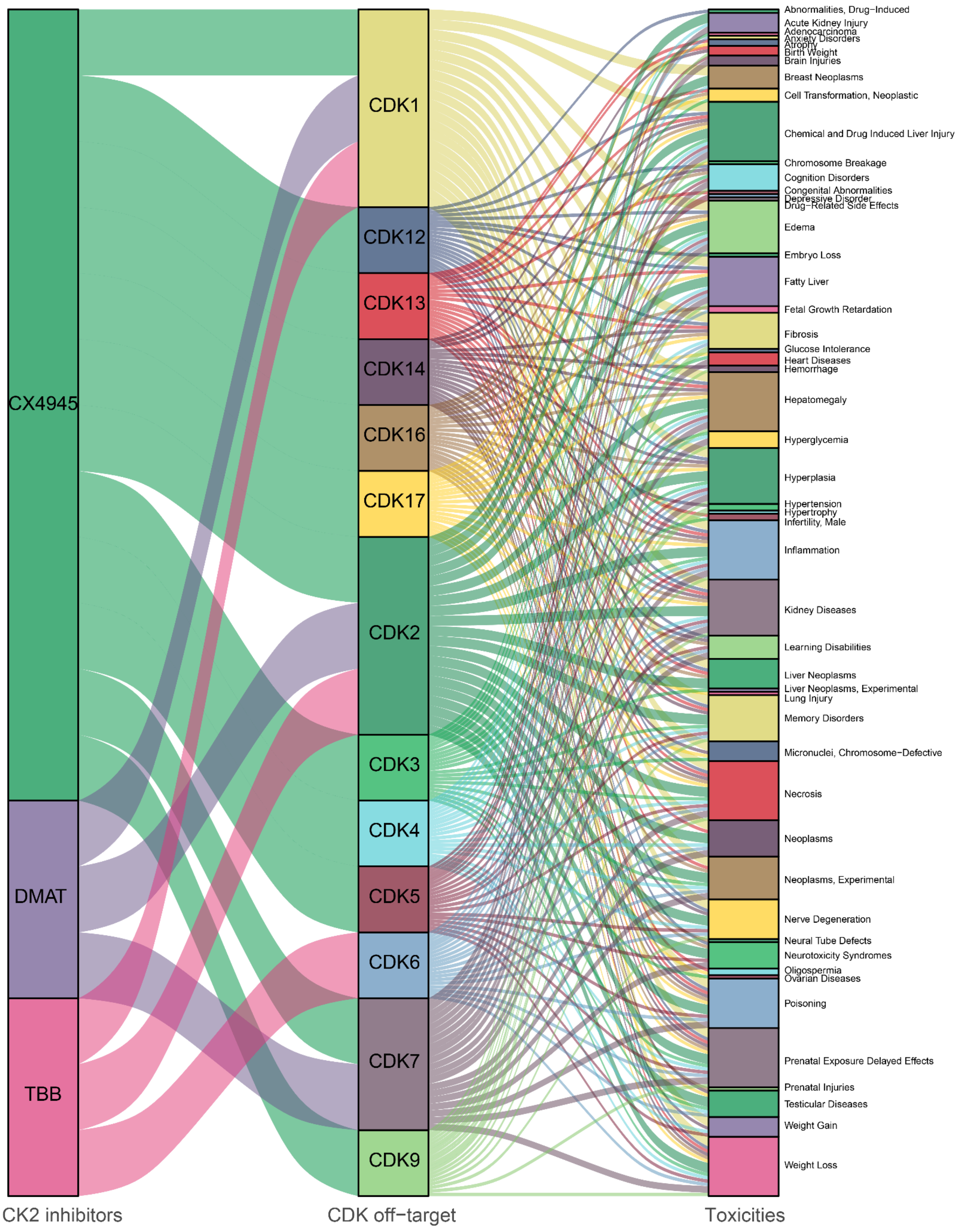


**Supplementary Figure 8.** Sankey diagram for the putative mechanistic links between CK2 inhibitors (CX4945, DMAT, TBB), CDK off-targets, and associated adverse effects.

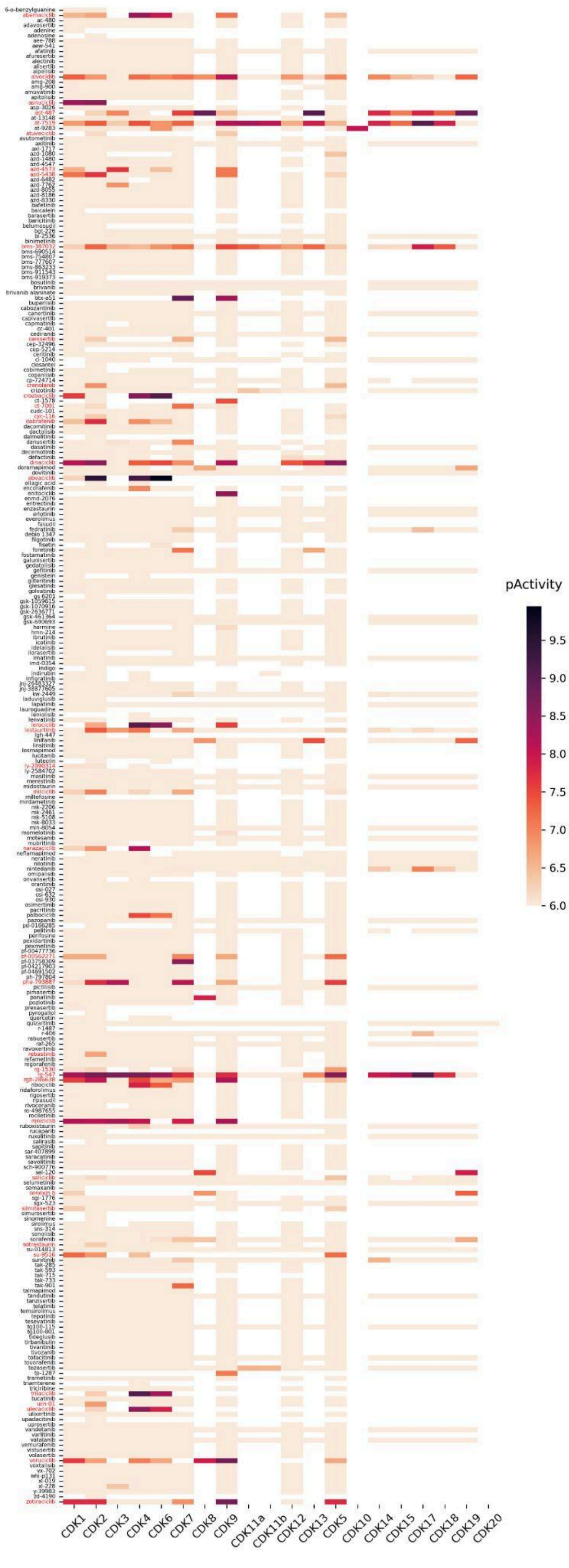


**Supplementary Figure 9**. **Publicly available compound bioactivity data for the CDK protein family.** The data was collected from ChEMBL, BindingDB, and DrugTargetCommons. Compound names colored in red have pActivity > 6 against CDK1 or CDK2. pActivity equals to -log10(molar IC50, EC50, Ki or Kd). A larger version of Supplementary Figure 9 can be found as a standalone file.

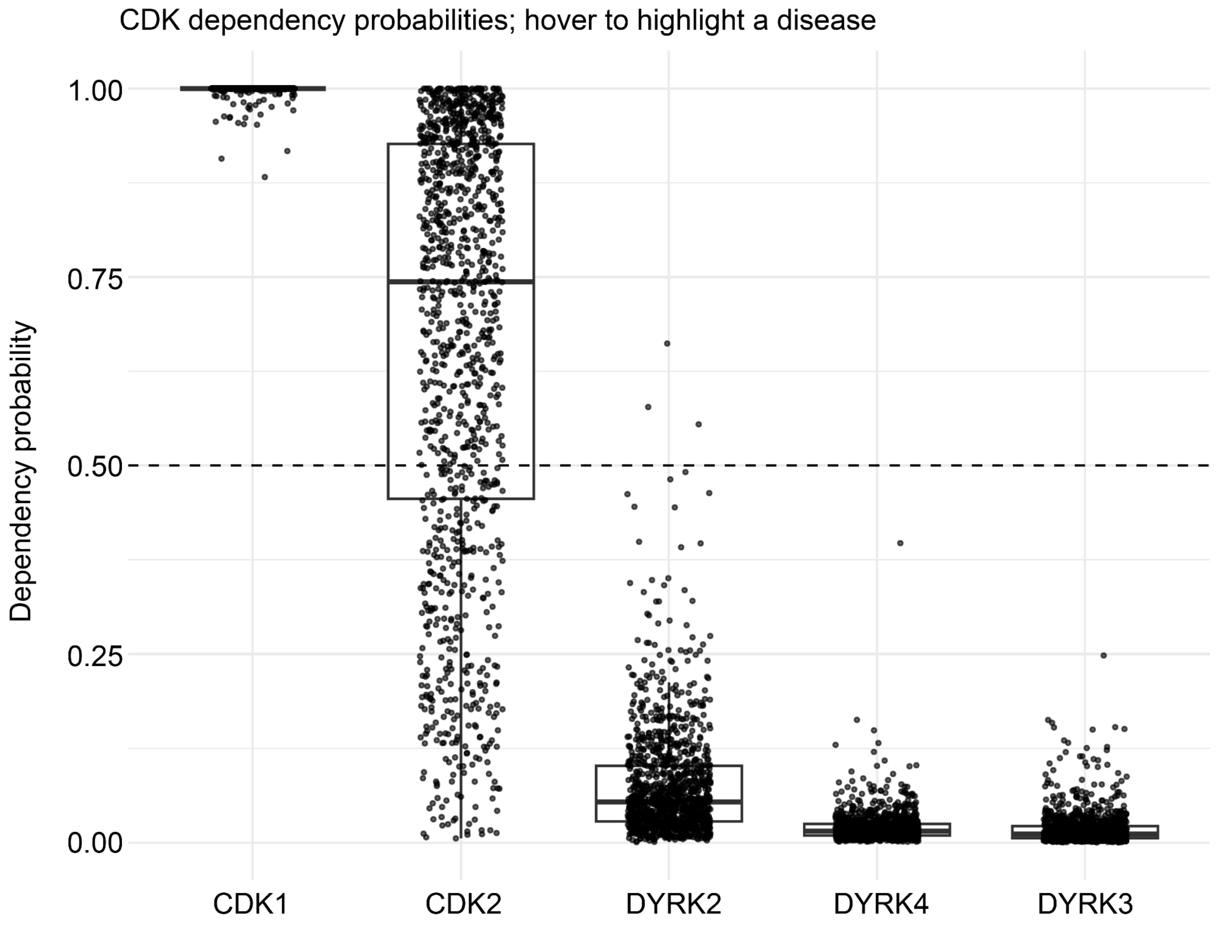


**Supplementary Figure 10**. **Cell line dependency probabilities for the selected kinases across cancer cell lines based on DepMap data.** CDK1 and CDK2 exhibit high dependency probabilities in a large proportion of cell lines, whereas DYRK family members (DYRK2, DYRK3, and DYRK4) show consistently low dependency, indicating minimal essentiality compared with CDKs.

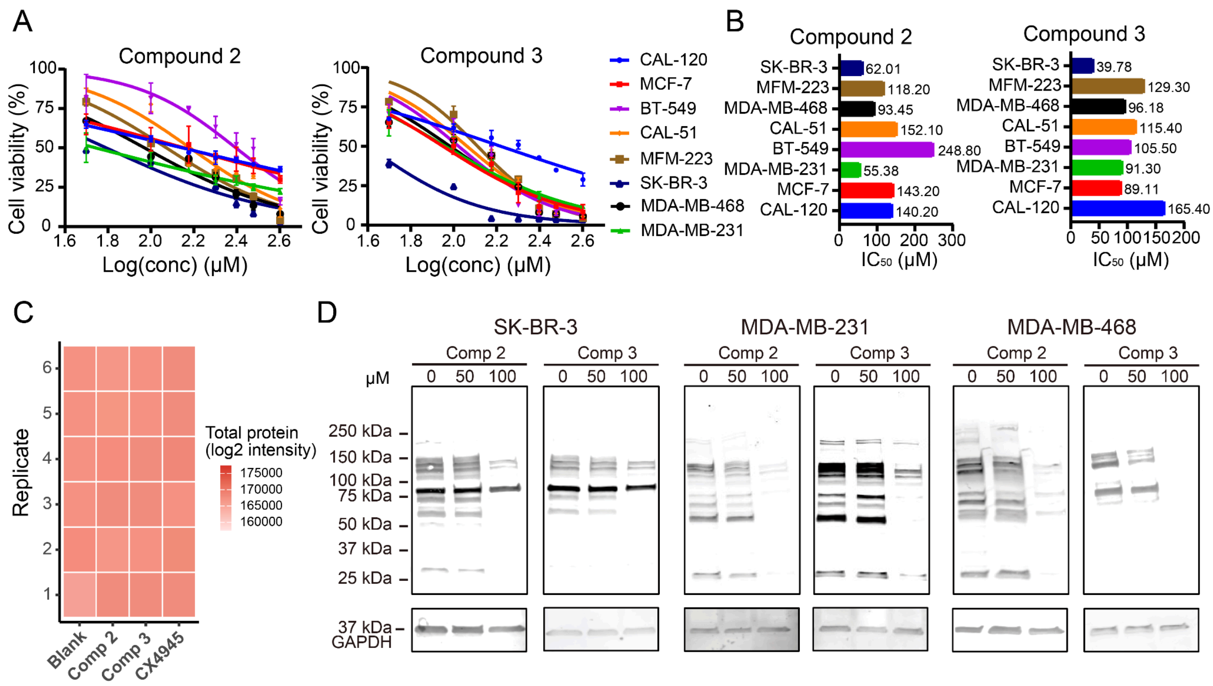


**Supplementary Figure 11**. **Cellular assays and specificity profiling.** (**A**) Inhibitory effects of the two active compounds (Compound 2 and Compound 3) on eight breast cancer cell lines, where CK2 is often overexpressed. (**B**) $IC_{50}$ values on the eight breast cancer cell lines were calculated by Graphpad Prism. (**C**) Heatmap of total protein expressions treated with different chemicals (comp2, comp3, CX4945) or no chemicals (Blank) in 6 independent replicates. (**D**) Immunoblot analysis of CK2-dependent substrate phosphorylation in SK-BR-3, MDA-MB-231, and MDA-MB-468 cells treated with Compound 2 (Comp 2) or Compound 3 (Comp 3). Cells were treated with the indicated concentrations (0, 50, and 100 μM), and CK2 substrate phosphorylation was detected using an anti-phospho CK2 substrate antibody. GAPDH was used as a loading control.

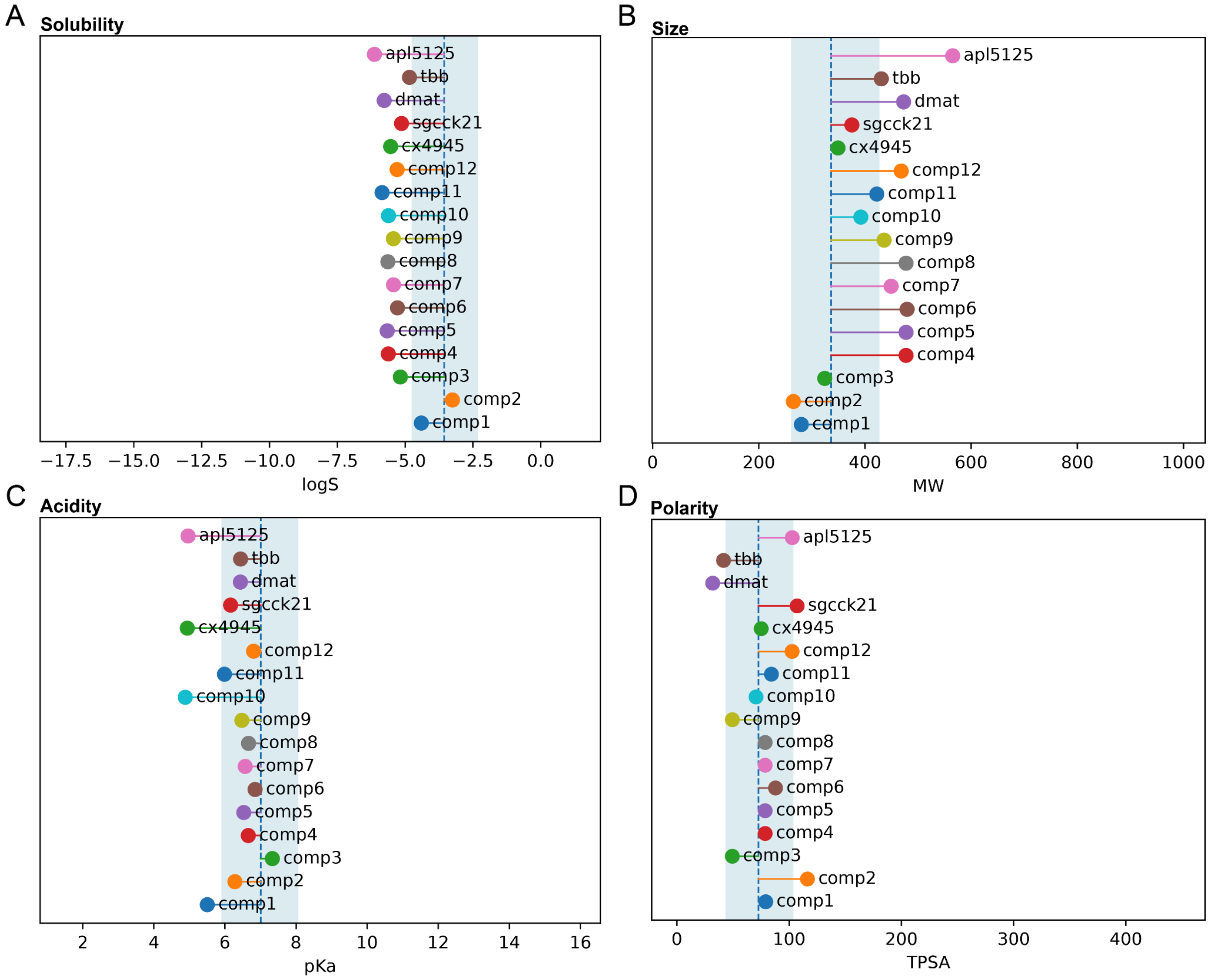


**Supplementary Figure 12. Predicted solubility, size, acidity and polarity of existing and novel CK2 inhibitors.** Different physicochemical properties (solubility, size, acidity, polarity) of the predicted molecules as well as control CK2 inhibitors. The blue color represents the interquartile range and the dashed line represents the median. The background data from which interquartile range was calculated is all of the small-molecule drugs present in the anatomical therapeutic chemical (ATC) classification system (Menestrina et al. 2025).
