## Supplementary figures and images for "Off-target mapping enhances selectivity of machine learning-predicted CK2 inhibitors"

### Large version of supplementary figure 9

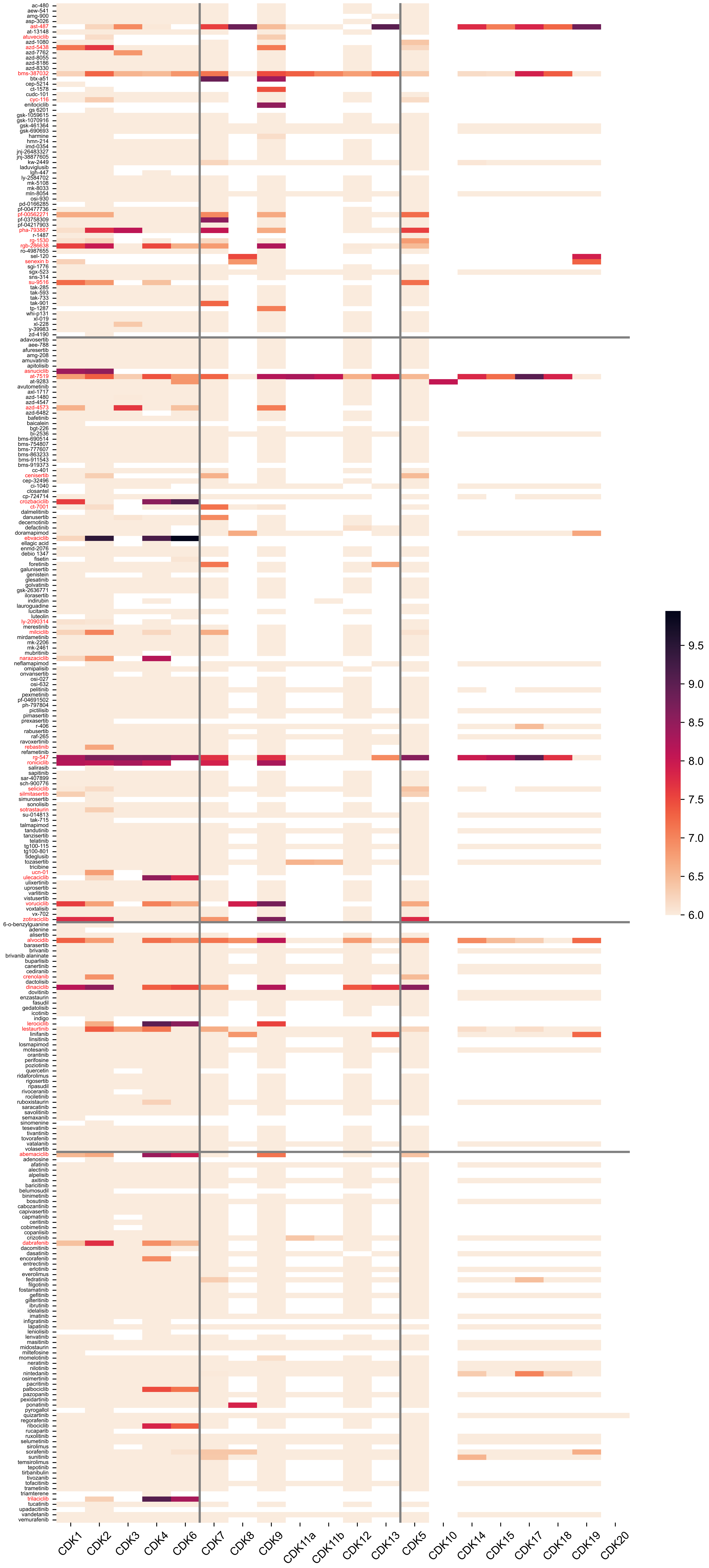
